# Grapevine holobiome metatranscriptomics provides a glimpse into the wood mycovirome

**DOI:** 10.1101/2025.03.21.644598

**Authors:** Humberto Debat, Marcos Paolinelli, Georgina Escoriaza, Sandra Garcia-Lampasona, Sebastián Gomez-Talquenca, Nicolás Bejerman

**Author notes:** Corresponding authors: Debat Humberto,; Nicolás Bejerman.

## Abstract

Given the agronomic and economic importance of viticulture, grapevine has been shown to host the largest number of viruses among plants to date. Nevertheless, studies assessing the grapevine-associated holobiont remain scarce. In this context, the viral component of this ecological niche is understudied. In this work, through metatranscriptomics of wood samples from individual grapevines that were either healthy or exhibited symptoms of grapevine trunk disease from Argentina, we provide a glimpse into the wood linked virome. Virus discovery from high-throughput sequencing data resulted in the identification and reconstruction of 123 novel virus sequences. Genetic and phylogenetic insights suggest that these sequences correspond to 78 novel virus species. Structural and functional annotation of the viruses showed a great diversity of genomic organizations, with the presence of dsRNA, ssRNA(-) and ssRNA(+) viruses belonging to more than 15 virus families. A significant number of viruses (66%) were linked to the recently accepted families *Botourmiaviridae, Narnaviridae* and *Mitoviridae*. Some highly divergent viruses resembling narnaviruses, ophioviruses, deltaflexiviruses and bunyaviruses could be accommodated within new genera or even new virus families. The differential detection and variable RNA levels across samples suggest complex dynamics and prevalence patterns of those novel viruses. The viral profile described here provides a first insight into the multifaceted South American grapevine wood holobiont mycovirome.

## 1. Introduction

Grapevine trunk diseases caused by trunk pathogens are becoming one of the main constraints on viticulture (Fontaine et al., 2016). These diseases are complex and often involve the infection of multiple fungal pathogens (Nerva et al., 2019A). Grapevine is gaining increasing significance on a global scale, prompting extensive exploration of its genetic variability and complexity (Grassi and de Lorenzis, 2021). A deeper understanding of this biodiversity emerges when viewing grapevine as a holobiont, an ecological unit comprising closely associated distinct species (Bettenfeld et al., 2022). This additional layer of biodiversity introduces a novel perspective on the etiology of diseases caused by microorganisms: the pathobiome, which encapsulates not only the pathogenic agent responsible for the disease but also other interacting microorganisms (Bass et al., 2019; Vayssier-Taussat et al., 2014), with grapevine trunk diseases serving as a clear example of this concept. The holobiome concept holds significance not only for providing a theoretical framework for pathobiome systems but also for emphasizing the role of symbionts in contributing to the phenotypic characteristics to an organism (Theis et al., 2016). Viruses, for instance, are often considered part of their host’s “enhanced genome” due to the diverse features they may confer (Bass et al., 2019).

Mycoviruses are viruses that can infect and replicate within fungi (Kondo et al., 2022) and have been found in a wide range of fungi (Chiapello et al., 2020; Garcia-Pedrajas et al., 2019; Ghabrial et al., 2015; Wu et al., 2025; Xie and Jiang, 2024; Lu et al., 2024). Most of them are cryptic, causing negligible detectable symptoms in the host (Jia et al., 2021). Despite their cryptic properties, viruses can potentially offer benefits beyond their asymptomatic nature. Certain pathogenic (acute) viruses exhibit intriguing biotechnological potential, particularly when their hosts are pests or pathogens. Mycoviruses, for instance, can be valuable in mitigating diseases caused by the phytopathogenic fungi they infect (Garcia-Pedrajas et al., 2019; Villain Larios et al., 2023; Xie and Jiang, 2024).

Numerous mycoviruses have been identified and characterized in the obligate biotrophic oomycete *Plasmopara viticola*, which colonizes grapevine (Chiapello et al., 2020), as well as in fungal pathogens of grapevine, such as *Neofusicoccum parvum* (Marais et al., 2022), *Botryosphaeriaceae* (Comont et al., 2024) and *Botrytis cinerea* (Ruiz-Padilla et al., 2021). Additionally, some mycoviruses have been found in endophytes within vine wood tissues (Nerva et al., 2019B). These findings illustrate the richness and diversity of the mycovirome in microorganisms associated with grapevine.

Here, through metatranscriptomics of wood samples from individual grapevines that were either healthy or exhibited symptoms of grapevine trunk disease in Argentina, we provide a glimpse into the wood mycovirome, expanding the knowledge of mycoviruses linked to grapevine fungal-associated diseases. We identified 78 new species, with the goal of enlightening the complexity of this pathosystem, favoring the integration of this information to foster the development of biotechnological platforms oriented to novel approaches for the biocontrol of this disease.

## 2. Material and Methods

### 2.1 Sample collection

Samples were collected as described by Paolinelli et al., (2022). Briefly, in Luján de Cuyo, Argentina, at the Estación Experimental Agropecuaria Mendoza INTA (−L33.005445,L−L68.864788), a vineyard housing ungrafted *Vitis vinifera* cv. Malbec plants, aged 23 years and trained in a vertical trellis system, underwent sampling at the onset of winter (09/11/2017). The plants belonged to clone 2 of the Malbec cultivar based on a previously characterized germplasm collection. Six plants were selected for analysis: three symptomatic for Hoja de malvón (HDM) disease (m11, m16, and m26) and three asymptomatic plants (c22, c23, and c25). These plants were uprooted and promptly transported to the laboratory, where they underwent surface sterilization with 70% ethanol. Subsequently, transversal cuts were made at the base and upper section of the trunk, as well as at both branch extremities. Woodchips obtained from bark, sapwood, and heartwood were collected and stored in sterile plastic bags and immediately frozen at −L80°C for later RNA extraction. Each plant sample comprised six subsamples, with the saw being sterilized using 70% ethanol (v/v) and 3% hydrogen peroxide (v/v) between each sampling event.

### 2.2. RNA extraction and Next generation sequencing

RNA was extracted and sequenced as described by Paolinelli et al., (2022). Briefly, the woodchip samples stored at −L80 °C were subjected to pulverization using liquid nitrogen in a mortar. Approximately 400 mg of the resulting pulverized plant tissues were employed for RNA extraction within 15-mL centrifuge tubes. The plant tissue underwent incubation with cetyl trimethyl ammonium bromide (CTAB) extraction buffer at 65 °C for 10 minutes. Subsequently, a chloroform-isopropyl alcohol mixture (24:1) was added, and the aqueous phase was collected post-centrifugation. Afterward, a precipitation step was carried out employing 3 M sodium acetate (pHL=L8). RNA was precipitated with 10 M lithium chloride, followed by two washing steps with 75% ethanol. To eliminate any DNA contamination, all obtained RNA underwent treatment with RQ1 RNase-Free DNase (Promega®, USA), and RNA was cleaned using the R1015 Zymo® kit as per the manufacturer’s instructions. Total RNA quantification was carried out using a Thermo Fisher® Nanodrop 2000, with quality assessment performed on a 1.5% agarose gel. Library preparation was performed using a TruSeq® Stranded Total RNA library preparation kit with Ribo-Zero™, and subsequent sequencing was conducted on an Illumina HiSeq4000® platform with 2L×L100 bp reads by Macrogen, Inc. (South Korea).

### 2.3 Identification of viruses

The trimming of TruSeq3-PE-2 adapters and the removal of reads with a Phred quality score below 30 was carried out using Trimmomatic v0.38 (Bolger et al., 2014), with the following parameters: “TruSeq3-PE-2.fa:2:30:10 SLIDINGWINDOW:4:15 LEADING:5 TRAILING:5 MINLEN:25”. Athough the libraries were constructed using Plant Ribo-Zero, remaining plant rRNA reads were identified through SortMeRNA v2.0 (Kopylova et al., 2012). To eliminate reads originating from the transcription of plastid and mitochondrial genomes, organelle sequences from the RefSeq database (https://www.ncbi.nlm.nih.gov/genome/organelle/), as well as DNA and RNA databases from plastids (ftp://ftp.ncbi.nlm.nih.gov/refseq/release/plastid/) and mitochondria (ftp://ftp.ncbi.nlm.nih.gov/refseq/release/mitochondrion/), were downloaded and used as a reference in Bowtie2 v2.3.4.3 (Langmed and Salzberg, 2012) mapping. The resultant filtered reads were then mapped onto the *Vitis vinifera* Pinot Noir genome (Jaillon et al., 2007) using STAR v2.6.1 (Dobin and Gingeras, 2015). Unmapped grapevine reads were extracted and *de novo* assembled using SPAdes version 3.11.1 (Bankevich et al., 2012). The assembled contigs were compared against the non-redundant protein database using the DIAMOND BLASTX program (version 0.9.10) with an E-value cutoff of 1 × 10^-5^ (Buchfink et al., 2015). Virus-like contigs were subsequently polished by re-mapping the filtered reads using Bowtie 2. The refined nucleotide viral sequences were imported into ORFinder, as implemented in https://www.ncbi.nlm.nih.gov/orffinder/, with a minimal 150 nt ORF length, and genetic code 4 parameters. Conserved domains of the predicted translated products were searched using the NCBI Conserved Domain Database v3.18 tool (CDD; www.ncbi.nlm.nih.gov/Structure/cdd/wrpsb.cgi).

### 2.4 Phylogenetic analysis

Multiple sequence alignments of RdRp sequences were carried out with the optimized automatic adjustment using MAFTT version 7 (Katoh et al., 2013) available at http://mafft.cbrc.jp/alignment/server/ (iterative refinement methods: L-INS-i strategy). The maximum-likelihood phylogenetic trees were then constructed using the MEGA11 (Tamura et al., 2021) with the WAG+G+I model.

## 3. Results and Discussion

### 3.1 Metatranscriptomic identification of mycoviruses from grapevine wood samples

Metatransciptomics analysis has greatly expanded our knowledge of the virosphere and has led to a dramatic increase in the number of mycoviruses discovered (Kotta-Loizou and Coutts 2017; Nerva et al. 2019; Chiapello et al. 2020; Nerva et al., 2019; Ruiz Padilla et al., 2021; Forgia et al., 2024; Lu et al., 2024; Wu et al., 2025; Liu et al., 2023A; Botella et al., 2022; Urzo et al., 2024; Marzano et al., 201). In our research, we analyzed six libraries from wood samples of individual grapevine plants, either healthy or showing symptoms of grapevine trunk disease from Argentina, which provides a glimpse of the wood virome. Virus discovery from the high-throughput sequencing data resulted in the assembly of 123 contigs, representing partial or coding-complete genome segments, which were assigned to 78 putative novel mycoviruses (Table 1). Most of the mycoviruses identified to date have RNA genomes, including positive single-stranded (+ss) RNA, negative ss (−ss) RNA, and double-stranded (ds) RNA (dsRNA) (Ayllon and Vainio, 2023; Kondo et al., 2022) and they have been classified into more than 23 families, although many remain unclassified (Hough et al., 2023). Structural and functional annotations of the 78 viruses identified in our study showed a great diversity of genomic organizations, with 65 putative mycoviruses harboring ssRNA (+) genomes, four containing dsRNA genomes, and the remaining nine having ssRNA (-) genomes (Table 1). The viruses discovered were assigned to more than 15 virus families, while some remain unclassified. Botourmia, narna-like, and mitoviruses were by far the most prevalent viruses identified (52/78), supporting other fungal virome studies that have identified botourmiaviruses, mitoviruses and narna-like viruses as the most abundant ones (Chiapello et al., 2020; Liu et al., 2023A; Jia et al., 2021; Jo et al., 2022; Ruiz-Padilla et al., 2021). Based on evolutionary clustering, some highly divergent viruses could be accommodated into new genera or even new virus families.

### 3.2 Molecular and phylogenetic characterization of the new identified viruses

#### 3.2A. Positive-sense single-stranded RNA viruses

##### Ambiguiviruses

Three contigs showed similarity to sequences from members of the proposed family “Ambiguiviridae”, which is a family of capsidless viruses with positive-sense ssRNA genomes that contains two ORFs in the same reading frame (Gilbert et al 2019). The three identified ambiguiviruses were named grapevine wood holobiome associated ambiguivirus 1 (GWHaAV1), grapevine wood holobiome associated ambiguivirus 2 (GWHaAV2), and grapevine wood holobiome associated ambiguivirus 3 (GWHaAV3). The coding-complete sequence of these viruses was obtained and deposited in the GenBank under accession numbers PQ474301, PQ474302, and PQ590144, respectively.

The coding-complete sequence of GWHaAV1 is 3214 nucleotides (nt) in length, while GWHaAV2 is composed of 3201 nt and GWHaAV3 sequence is 4096 (Table 1). All three viruses contain two open reading frames (ORF1 and ORF2) (Fig. 1A), which are in the same reading frame, thus they are likely expressed as a fusion protein by the readthrough of the UAG stop codon. ORF1 of GWHaAV1, GWHaAV2 and GWHaAV3 encodes a hypothetical protein (HP) of 247 amino acids (aa), 234 aa, and 320 aa, respectively. Notably, GWHaAV2 HP contains a highly conserved arginine-rich motif (RxRRR), which, in the tombusviruses, is critical for efficient RNA binding (Jia et al., 2023). BLASTp search against the nr protein database revealed that the best matches for the HP of GWHaAV1, GWHaAV2, and GWHaAV3 were proteins coded by rice tombus-like virus 3 (coverage, 74%; E-value, 6e-30; identity, 44.27%), Erysiphe necator associated ambiguivirus 2 (coverage, 70%; E-value, 8e-26; identity, 40.61%), and soybean leaf associated ssRNA virus 1 (coverage, 61%; E-value, 4e-33; identity, 42.08%), respectively. ORF2 of GWHaAV1, GWHaAV2 and GWHaAV3 encodes an RNA-dependent RNA polymerase (RdRp) RdRp of 498, 481 aa, and 526 aa, respectively, sharing less than 50% identity between them. The RdRp sequences exhibit similarity to those of members of the proposed “*Ambiguiviridae*” family; whose name reflects the unusual GDN triad within the RdRp motif and the uncertainty regarding the host of Diaporthe RNA Virus, the first virus described in this group (Gilbert et al., 2019). The seven core RdRp motifs present in viral RdRps (Jia et al., 2023) were also identified in the GWHaAV1, GWHaAV2 and GWHaAV3 RdRp’s. Like other reported ambiguiviruses (Gilbert et al., 2019; Zhong et al., 2022; Jia et al., 2023; Buivydaite et al., 2024), a GDN tripeptide was identified in motif C of GWHaAV1, GWHaAV2, and GWHaAV3, replacing the highly conserved GDD motif generally found in (+) ssRNA viruses. It has been demonstrated that mutating GDD to GDN reduces the catalytic activity of RdRp in (+) ssRNA viruses (Vasquez et al., 2000). A BLASTp search of the nr protein database showed that the closest matches for the RdRp of GWHaAV1, GWHaAV2 and GWHaAV3 were the RdRp of Alternaria dianthicola umbra-like virus 1 (coverage, 99%; E-value, 5e-173; identity, 55.82%, and coverage, 97%; E-value, 4e-162; identity, 54.97%, respectively) and grapevine associated tombus-like virus 4 (coverage, 3%; E-value, 0.0; identity, 54.77%).

**Figure 1.**
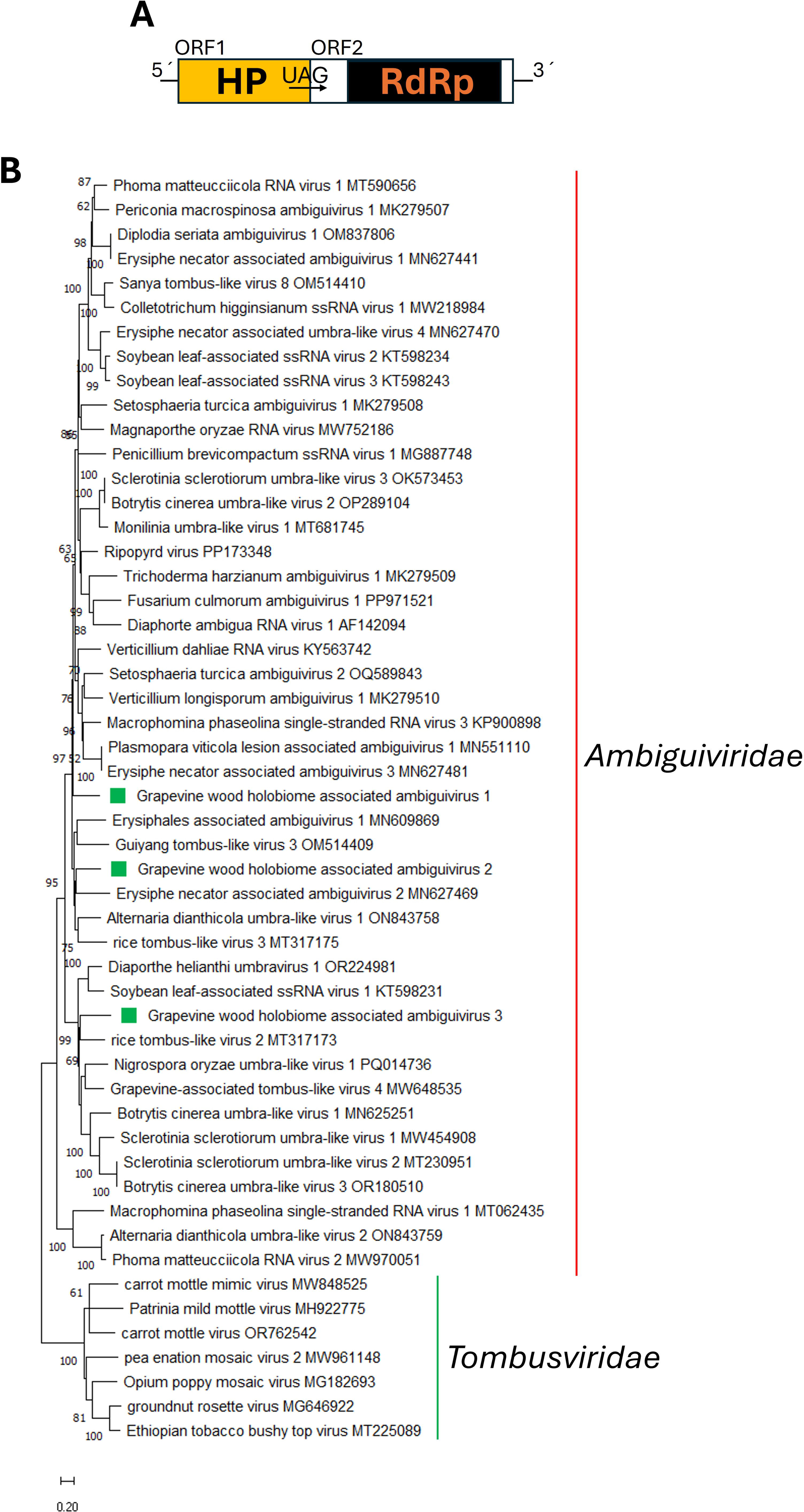
**(A)** Schematic representation of the genome organization of grapevine wood holobiome associated ambiguivirus 1-3. **(B)** Maximum likelihood phylogenetic trees reconstructed using the RdRp protein sequence of grapevine wood holobiome associated ambiguivirus 1-3 and of representative ambiguiviruses. Tombusvirids sequences were used as outgroup. Bootstrap values above 50% are shown (1000 replicates). The grapevine wood holobiome associated ambiguivirus 1-3 are indicated with a green square. The scale bar shows the substitution per site.

A fusion protein is expressed by GWHaAV1, GWHaAV2 and GWHaAV3 ORF’s 1 and 2. Similar to most of the ambiguiviruses reported so far (Gilbert et al., 2019; Jia et al., 2023; Buivydaite et al., 2024), the translation termination codon of GWHaAV1, GWHaAV2 and GWHaAV3 is a UAG (amber). A similar P1-P2 readthrough protein is also encoded by members of the plant virus family *Tombusviridae*, which is closely related to the proposed “*Ambiguivirdae*” family, which is composed of fungi-associated members. Nevertheless, unlike the tombusviruses, ambiguiviruses do not encode a movement protein (MP) or a capsid protein (CP). It has been hypothesized that, considering the close relationship between plants and fungi and the host adaptation of plant-infecting viruses, ambiguiviruses and tombusviruses may share a common ancestral origin but diverged during adaptation to their respective hosts (Jia et al., 2023).

A phylogenetic tree based on RdRp aa sequence alignment placed GWHaAV1, GWHaAV2 and GWHaAV3 within distinct clusters of ambiguiviruses (Fig. 2B). GWHaAV1 formed a monophyletic clade clustering with other proposed ambiguiviruses, while GWHaAV2 clustered separately with Erysiphe necator associated ambiguivirus 2 (Fig. 2B), and GWHaAV3 clustered with rice tombus-like virus 2 (Fig. 2B). Thus, based on their RdRp identity values, GWHaAV1, GWHaAV2 and GWHaAV3 represent three novel members of the proposed “*Ambiguiviridae*” family. GWHaAV1and GWHaAV2 should be classified into one genus, while GWHaAV3 belongs to a separate genus, both of which require formal taxonomic designation. Moreover, the demarcation criteria based on RdRp protein identity still need to be established to define species boundaries within each proposed genus of the proposed set to accommodate species between each proposed genus within the “*Ambiguivirdae*” family.

**Figure 2.**
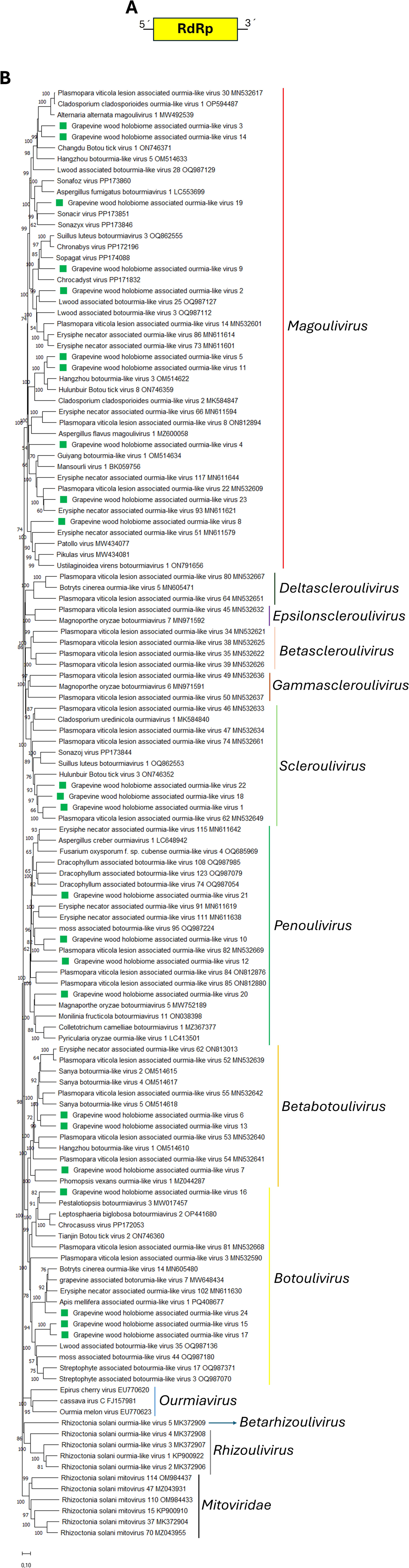
**(A)** Schematic representation of the genome organization of grapevine wood holobiome associated ourmia-like virus 1-24. **(B)** Maximum likelihood phylogenetic trees reconstructed using the RdRp protein sequence of grapevine wood holobiome associated ourmia-like virus 1-24 and of representative botourmiaviruses. Mitovirus sequences were used as outgroup. Bootstrap values above 50% are shown (1000 replicates). The grapevine wood holobiome associated ourmia-like virus 1-24 are indicated with a green square. The scale bar shows the substitution per site.

##### Botourmiaviruses

Twenty-four contigs showed similarity to sequences from members of the family *Botourmiaviridae*, a family of positive-sense ssRNA viruses that infect plants and fungi. Plant-associated members have a tripartite genome, while fungi-associated botourmiaviruses are non-encapsidated, with a monopartite and monocistronic genome (Ayllon et al., 2020). The *Botourmiaviridae* family comprises twelve genera, of which one (*Ourmiavirus*) consists of plant-associated members, while the other eleven contain fungi-associated members (Ayllon et al., 2020; https://ictv.global/taxonomy). Among fungi-associated boutormiavirids, genome length ranges from 2100 to 5185 nt, and all contain a single ORF encoding an RdRp (Wang et al., 2022). The identified botourmiaviruses were named grapevine wood holobiome-associated ourmia-like virus 1-24 (GWHaOLV1-24). The full-length sequence of fourteen of these twenty-four viruses were assembled and deposited in GenBank under accession numbers PQ773447, PQ773448, PQ773449, PQ773450, PQ773451, PQ773452, PQ773455, PQ773457, PQ773458, PQ773460, PQ773466, PQ773467, PQ773469 and PQ773470 (Table 1). The remaining ten viruses GWHaOLV7, GWHaOLV8, GWHaOLV10, GWHaOLV13, GWHaOLV15, GWHaOLV16, GWHaOLV17, GWHaOLV18, GWHaOLV19, and GWHaOLV22 were partially assembled and deposited under accession numbers PQ773453, PQ773454, PQ773456, PQ773459, PQ773461, PQ773462, PQ773463, PQ773464, PQ773465 and PQ773468 (Table 1). Among the fourteen viruses where the full-length coding region was assembled, genome size ranged between 2351 nt and 3011 nt (Table 1). Like all fungi-associated botourmiaviruses, all twenty-four viruses contain a single ORF encoding a putative RdRp (Fig. 2A). BLASTp searches using the deduced RdRP aa sequence revealed that the closest matches for GWHaOLV1-24 were Plasmopara viticola lesion associated ourmia-like virus 62, Lwood associated botourmia-like virus 25, Hangzhou botourmia-like virus 5, Guiyang botourmia-like virus 1, Hulunbuir Botou tick virus 8, Erysiphe necator associated ourmia-like virus 62, Phomopsis vexans ourmia-like virus 1, Yellow silver pine associated botourmia-like virus 66, Sopagat virus, Plasmopara viticola lesion associated ourmia-like virus 82, Hangzhou botourmia-like virus 3, Botourmiviridae sp, Sanya botourmia-like virus 4, Hangzhou botourmia-like virus 5, Streptophyte associated botourmia-like virus 17, Pestalotiopsis botourmiavirus 3, Streptophyte associated botourmia-like virus 17, Sopafab virus, Plasmopara viticola lesion associated ourmia-like virus 62, Sonacir virus, Colletotrichum camelliae botourmiavirus 1, Botourmiviridae sp, Plasmopara viticola lesion associated ourmia-like virus 62, Plasmopara viticola lesion associated ourmia-like virus 22 and Botrytis cinerea ourmia-like virus 14 RdRp’s, respectively (Table 1), with identities values below 90%. Moreover, the aa identities among those viruses assembled in this study ranged between 22.95% to 83.2%.

Phylogenetic analysis based on RdRp aa sequence alignment revealed that GWHaOLV 1-24 clustered into five well-supported clades (Fig. 2B). Specifically, ten viruses (GWHaOLV2-5, GWHaOLV8-9, GWHaOLV11, GWHaOLV14, GWHaOLV19 and GWHaOLV23) clustered with members of the genus *Magoulivirus*, three viruses (GWHaOLV1, GWHaOLV18 and GWHaOLV22) clustered with members of the genus *Scleroulivirus*, four viruses (GWHaOLV10, GWHaOLV12 and GWHaOLV20-21) clustered with members of the genus *Penoulivirus*, three viruses (GWHaOLV6-7, and GWHaOLV13) clustered with members of the genus *Betabotoulivirus*, and four viruses (GWHaOLV15-17 and GWHaOLV24) clustered with members of the genus *Botoulivirus* (Fig. 9B). According to the ICTV species demarcation criteria, classification is based on sequence identity in the complete RdRp (less than 90 % aa sequence identity) (Ayllon et al., 2020). Therefore, the fourteen botourmiavirids whose complete RdRp sequences were assembled represent novel members of the family *Botourmiaviridae*. Within the family, GWHaOLV2-5, GWHaOLV9, GWHaOLV11, GWHaOLV14 and GWHaOLV23 are novel magouliviruses, GWHaOLV1 is a novel scleroulivirus, GWHaOLV12 and GWHaOLV20-21 are novel penouliviruses, GWHaOLV6 is a novel betabotoulivirus and GWHaOLV24 is a novel botoulivirus. Therefore, this study revealed significant botourmiavirus diversity, as the newly discovered viruses can be classified into five distinct genera

Many ourmia-like viruses were identified in plant-pathogenic fungi in recent years (Chiapello et al., 2020; Li et al., 2021; Xie and Jiang, 2024). In-depth studies of some of these viruses have demonstrated that they can replicate and move between fungal strains via hyphal anastomosis, suggesting that the RdRp alone is sufficient for replication, infection, and transmission of ourmi-like viruses in their fungal hosts (Wang et al., 2020). These findings support a hypothesis of ourmia-like virus evolution, where these viruses may have evolved from plant-associated ourmiaviruses trough gene-loss events during their adaptation to fungal hosts (Marzano et al., 2016; Donaire et al., 2016). Thus, the identification of the twenty-four ourmia-like viruses in this study contributes to a deeper understanding of the taxonomy and evolution of mycoviruses, highlighting their potential host shifts and genomic adaptations over time.

##### Deltaflexiviruses

Three contigs showed similarity to sequences from members of the family *Deltaflexiviridae*, which includes a single genus, *Deltaflexivirus* (Wu et al., 2024). Its members have a positive-sense ssRNA genomes that can be segmented (Mu et al., 2021) or unsegmented (Xiao et al., 2023, Li et al., 2016; Chen et al., 2016; Hamid et al., 2018), with the gene encoding the replicase being the only one conserved among all deltaflexiviruses (Xiao et al., 2023). The three identified deltaflexiviruses were named grapevine wood holobiome associated deltaflexivirus 1-3 (GWHaDFV1-3). The coding-complete sequence of GWHaDFV1, as well as the partial sequences of GWHaDFV2 and GWHaDFV3, were obtained and deposited in GenBank under accession numbers PQ567243, PQ567244 and PQ567245, respectively.

The coding-complete sequence of GWHaDFV1 is 8592 nt in length and contains one large ORF (ORF1) and four small downstream ORFs (ORF2-5) (Table 1, Fig. 3A). This genomic organization is similar to that of Fusarium graminearum deltaflexivirus 1 (FgDFV1) (Chen et al., 2016), Fusarium deltaflexivirus 2 (FDFV2) (Wu et al., 2024), and Calypogeia fissa associated deltaflexivirus (CafADV) (Mifsud et al., 2022). In contrast, four ORFs were predicted in GWHaDFV2, while only one ORF was identified in GWHaDFV3. ORF1 encodes a putative replication-associated polyprotein containing three conserved domains: Mtr (methytransferase), Hel (helicase), and RdRp, which are present in all members of the family *Deltaflexiviridae* (Xiao et al., 2023). The other ORFs encode hypothetical proteins without conserved domains. Moreover, deltaflexiviruses are thought to be capsidless, suggesting an intracellular mode of transmission (Xiao et al., 2023). BLASTp searches using the deduced aa sequence showed that the best hit for GWHaDFV1 complete replication-associated polyprotein (1988 aa) and GWHaDFV2 and GWHaDFV3 partial replication-associated polyproteins, was FDFV2 (coverage, 97%; E-value, 0.0; identity 58.51% for GWHaDFV1), Erysiphe necator associated deltaflexivirus 3 and Pestalotiopsis deltaflexivirus 1 replication-associated polyprotein, respectively (Table 1). According to ICTV species demarcation criteria, classification is based on host range, number of minor ORFs, and sequence identity in the replication-associated polyprotein (less than 70% aa sequence identity) (Xiao et al., 2023). Therefore, GWHaDFV1should be classified as novel species within the *Deltaflexiviridae* family.

**Figure 3.**
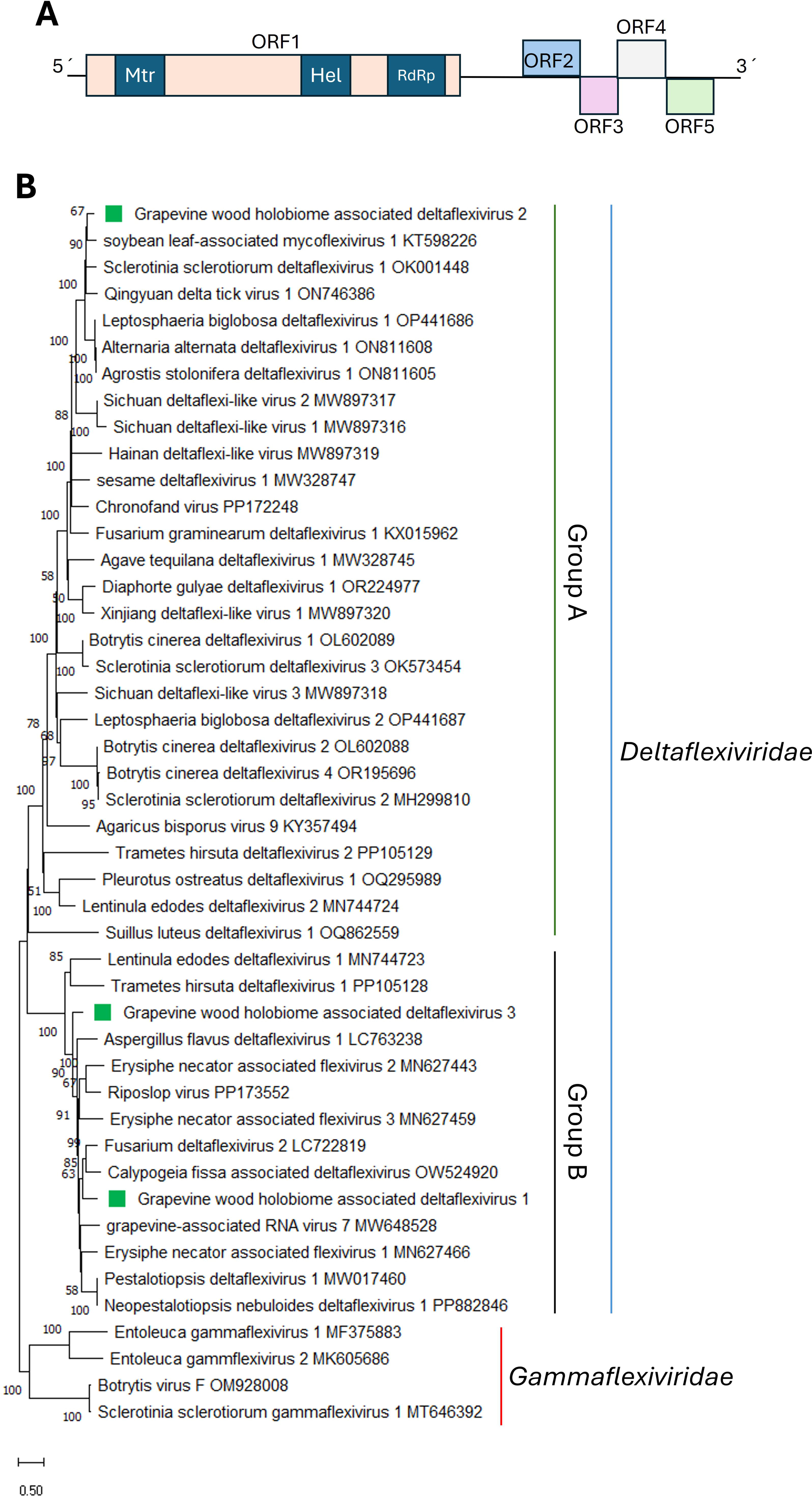
**(A)** Schematic representation of the genome organization of grapevine wood holobiome associated deltaflexivirus 1. **(B)** Maximum likelihood phylogenetic trees reconstructed using the replication-associated polyprotein sequence of grapevine wood holobiome associated deltaflexivirus 1-3 and of representative deltaflexiviruses. Gammaflexivirus sequences were used as outgroup. Bootstrap values above 50% are shown (1000 replicates). The grapevine wood holobiome associated deltaflexivirus 1-3 are indicated with a green square. The scale bar shows the substitution per site.

The recent discovery of the novel virus Fusarium oxysporum icosahedral virus 1 (FoIV1), referred as Fusarium deltaflexivirus 2 (FDFV2) in the NCBI nucleotide core database, suggests that deltaflexivirus particles are likely icosahedral, similar to tymoviruses, rather than filamentous, like the Alpha-, Beta-and Gammaflexiviruses (Wu et al., 2024). FolV1 ORF4 encodes a single jelly-roll (SJR)-like coat protein (CP), and clustering analysis suggested that FoIV1-related viruses have virions composed of SJR-like CPs, rather than being capsidless, as observed in FoIV1 (Wu et al., 2024). When the protein encoded by GWHaDFV1 ORF4 was analyzed using HHpred, significant hits to the CP of tymoviruses were obtained, supporting the hypothesis that GWHaDFV1 ORF4 also encodes a putative CP. It has been hypothesized that the ancestral virus of the deltaflexivirids used plants as its primordial host, and some of its members, such as Sclerotinia sclerotorium deltaflexivirus 2 and 3 and Lentinula edodes deltaflexivirus 1, lost their CP during evolution (Wu et al., 2024).

Phylogenetic analysis based on the replication-associated polyprotein aa sequence alignment revealed that GWHaDFV1, GWHaDFV2 and GWHaDFV3 formed a well-supported clade with other deltaflexiviruses, where two groups, previously named “group A” and “group B” (Wu et al., 2024), were observed (Fig. 3B). Most members of “group A” have genomes composed of four ORFs, whereas most members of “group B” have genomes composed of five ORFs (Wu et al., 2024). GWHaDFV1 and GWHaDFV3 were placed within “group B” (Fig. 3B), while GWHaDFV2 was placed within “group A” clustering with soybean leaf associated mycoflexivirus 1 (Fig. 3B), which has a similar genomic organization (Marzano et al., 2016). Multiple alignment of the deltaflexivirids RdRp domain showed that GWHaDFV1 and other “Group B” viruses share canonical RdRp motifs A-C (Urayama et al., 2024), retaining acidic residues similar to those reported for FolV1 and other “Group B” members (Wu et al., 2024). In contrast, GWHaDFV2 and members of “Group A” exhibit divergent RdRp motifs (Wu et al., 2024). Moreover, GWHaDFV1 has five ORFs, confirming its placement with “Group B”, while GWHaDFV2 has four ORFs, confirming its phylogenetic placement with “Group A” (Wu et al., 2024). These results suggest that viruses grouped within “Group A” and “Group B” should be classified into two new genera within the family *Tymoviridae*, without the term “flexiviridae”, or possibly within a new family in the order *Tymovirales*, as suggested by Wu et al., (2024). Accordingly, GWHaDFV1 represents a novel member of the newly assigned genus/family along with other “Group B” members, whereas GWHaDFV2 represents a novel member of the newly assigned genus/family with “Group A” members.

##### Hypovirus

One contig showed similarity to sequences from members of the family *Hypoviridae*, which is a family of capsidless viruses comprising eight genera. Its members have mono-or bicistronic positive sense genomes, with one long ORF or two ORFs (Chiba et al., 2023). The putative hypovirus was named grapevine wood holobiome associated hypovirus (GWHaHV). The coding-complete sequence of this virus was obtained and deposited in GenBank under accession number PQ673554.

The coding-complete sequence of GWHaHV is 13686 nt in length (Table 1) and contains one ORF, which encodes a putative replication-associated polyprotein (Fig. 4A). This genomic organization is similar to that of Alternaria alternata hypovirus 1 (AaHV1) (Li et al., 2019), Fusarium sacchari hypovirus 1 (FsHV1) (Yao et al., 2020) and Beticola hypovirus 1 (BHV1) (Li et al., 2021), among other hypovirues. The encoded replication-associated protein is 4141 aa in length, and three conserved domains were identified in its sequence: DUF3525, RNA_pol_sf, and HEL, located at aa positions 2264-2608, 2743-3198 and 3643-3788, respectively. Moreover, GWHaHV replication-associated protein has a conserved SDD tripeptide in the RdRP motif, which was also found in the replication-associated polyproteins of AaHV1 and FsHV1 (Yao et al., 2020). Like AaHV1 (Li et al., 2019), a papain-like cysteine protease motif was not detected in the GWHaHV replication-associated protein. Interestingly, alignment of the N-terminal region of the GWHaHV protein with the papain-like cysteine protease domain regions of other hypo- and hypo-like viruses failed to reveal the presence of the three conserved cysteine protease core residues (cysteine, histidine, and glycine) (Koonin et al., 1991), which have been identified in other hypo-and hypo-like viruses (Li et al., 2019). Thus, it remains unknown whether GWHaHV-encoded polyprotein is processed by cysteine proteases. The results of BLASTp searches of the nr protein database showed that the best hit was the replication-associated polyprotein encoded by BHV1 (coverage, 84%; E-value, 0.0; identity, 56.26%).

**Figure 4.**
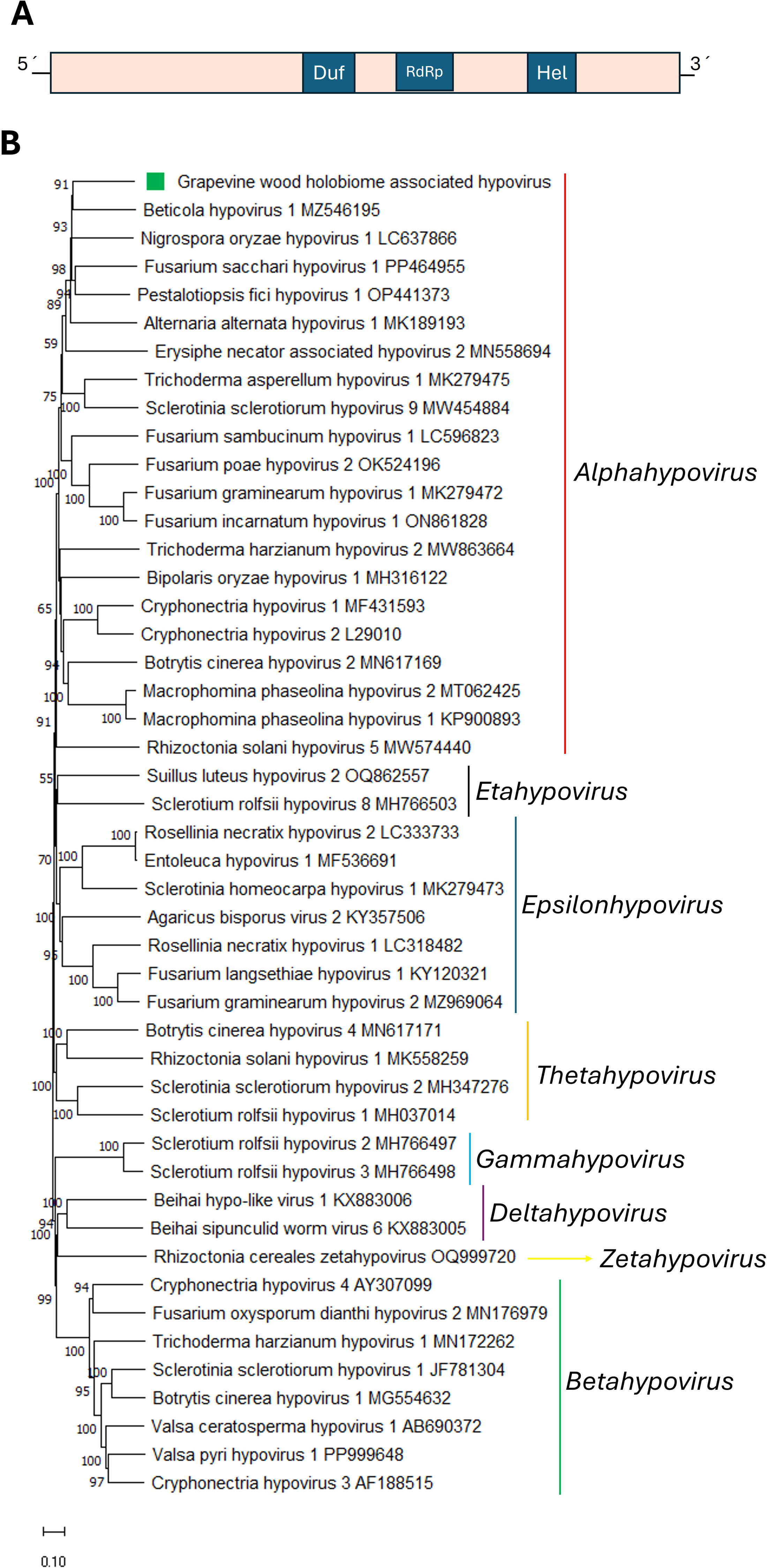
**(A)** Schematic representation of the genome organization of grapevine wood holobiome associated hypovirus. **(B)** Maximum likelihood phylogenetic trees reconstructed using the replication-associated polyprotein sequence of grapevine wood holobiome associated hypovirus and of representative hypoviruses. Bootstrap values above 50% are shown (1000 replicates). The grapevine wood holobiome associated hypovirus is indicated with a green square. The scale bar shows the substitution per site.

A phylogenetic tree based on replication-associated polyprotein aa sequence alignment placed GWHaHV in a well-supported clade with other alphahypoviruses (Fig. 4B), where it clustered with BHV1. According to ICTV species demarcation criteria, classification is based on sequence identity in the complete polyprotein (<80% aa identity). Therefore, GWHaHV1 is a novel member of the *Alphahypovirus* genus within the family *Hypoviridae*.

Several hypoviruses have been reported to induce hypovirulence to their fungal host, making them potential biocontrol tools for sustainable plant disease management (Garcia-Pedrajas et al., 2019; Li et al., 2019). Therefore, further studies on the molecular characterization and the pathogenicity of GWHaHV should be conducted to evaluate whether this virus could be used as a biocontrol agent for fungal crop disease management.

##### Mitoviruses

Six contigs showed similarity to sequences from members of the family *Mitoviridae*. which is characterized by capsidless viruses with positive-sense ssRNA genomes containing a single ORF (Sadiq et al., 2022; Wolf et al., 2018), and is composed of four genera (Jacquat et al., 2023). The six identified mitoviruses were named grapevine wood holobiome associated mitovirus 1-6 (GWHaMV1-6). The partial sequence of these six viruses were obtained and deposited in GenBank under accession numbers PQ729981, PQ729982, PQ729983, PQ729984, PQ729985 and PQ729986, respectively (Table 1).

All six viruses contain a single ORF encoding a putative RdRp (Fig. 5A), and the Mitovir_RNA_pol (Mitovirus RNA-dependent RNA polymerase; pfam05919) conserved motif was identified in all encoded RdRp’s. Moreover, the RdRp catalytic motif GDD (Charon et al., 2022) was also found in the RdRp of all six identified mitoviruses. BLASTp searches using the deduced aa sequences showed that the best hit for GWHaMV1, GWHaMV2, GWHaMV3, GWHaMV4, GWHaMV5 and GWHaMV6 was Rhizoctonia solani mitovirus 15, Hainan mito-like virus 26, Rhizoctonia cerealis duamitovirus, grapevine-associated mitovirus 2, Sichuan mountain mitovirus 12 and Guangxi riberbank mitovirus 3, respectively (Table 1). Phylogenetic analysis based on RdRp aa sequence alignment revealed that GWHaMV1, GWHaMV2, GWHaMV3, GWHaMV4, GWHaMV5 and GWHaMV6 were placed in a well-supported clade with other mitoviruses classified in the genus *Duamitovirus* (Fig. 5B). Specifically, GWHaMV1 clustered with Rhizoctonia solani mitovirus 47, 88 and 114, and grapevine-associated mitovirus 10 (Fig. 5B), GWHaMV2 clustered with Hainan mito-like virus 26 (Fig. 5B), GWHaMV3 clustered with Rhizoctonia solani mitovirus 11, 58 and 96 (Fig. 5B), GWHaMV4 with grapevine-associated mitovirus 2 (Fig. 5B), GWHaMV5 clustered with Sichuan mountain mitovirus 12 (Fig. 5B), while GWHaMV6 clustered separately with Guangxi riberbank mitovirus 3 (Fig. 5B).

**Figure 5.**
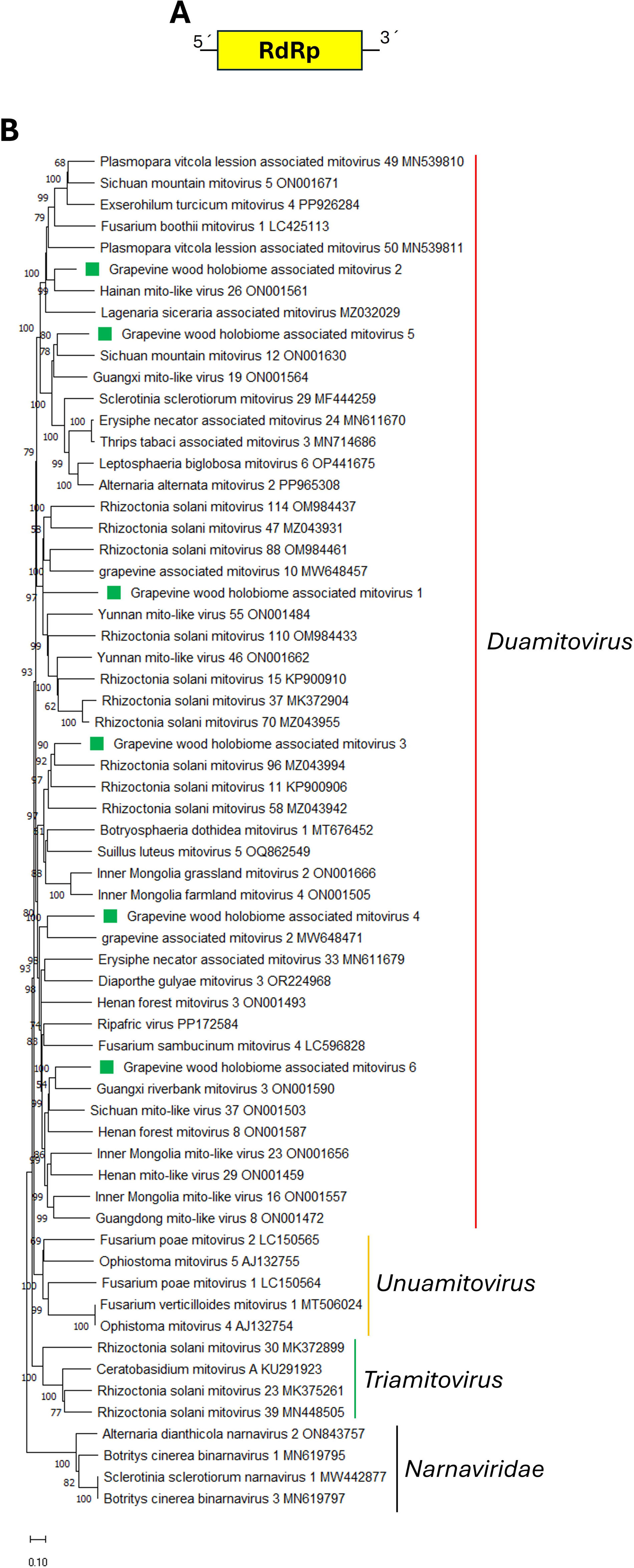
**(A)** Schematic representation of the genome organization of grapevine wood holobiome associated mitovirus 1-6. **(B)** Maximum likelihood phylogenetic trees reconstructed using the RdRp protein sequence of grapevine wood holobiome associated mitovirus 1-6 and of representative mitoviruses. Narnavirus sequences were used as outgroup. Bootstrap values above 50% are shown (1000 replicates). The grapevine wood holobiome associated mitovirus 1-6 are indicated with a green square. The scale bar shows the substitution per site.

##### Narna-like viruses

###### Monosegmented

Five contigs showed similarity to monosegmented narna-like virus sequences. *Narnaviridae* is a family of capsidless viruses with positive-sense ssRNA genomes containing a single ORF (Hillman and Cai, 2013) that belong to the order *Wolframvirales*. Currently, only two species have been accepted by the ICTV and classified within the genus *Narnavirus* (https://ictv.global/taxonomy). Therefore, most of the narna-like viruses reported to date, which exceed a thousand, remain unclassified. Moreover, narna-like viruses assembled in metagenomic studies may contain additional genes and multiple ORFS, including a capsid protein (Sadiq et al., 2022). Some of these more complex narna-like viruses have been proposed for classification within the family “*Narliviridae*” in the order *Ourlivirales*, which also includes the family *Botourmiaviridae* (Sadiq et al., 2022). The five identified monosegmented narna-like viruses were named as grapevine wood holobiome associated narna-like virus 1-5 (GWHaNLV1-5). The full length-sequences of GWHaNLV1-4 were assembled and deposited in GenBank under accession numbers PV021966, PV021967, PV021968 and PV021969 (Table 1). For GWHaNLV5, only a partial sequence was assembled and deposited in GenBank under accession number PV021970 (Table 1). The four viruses with fully assembled coding-sequences contain a single ORF encoding an RdRp (Fig. 6A), with their genome lengths ranging between 2488 nt and 3383 nt (Table 1).

**Figure 6.**
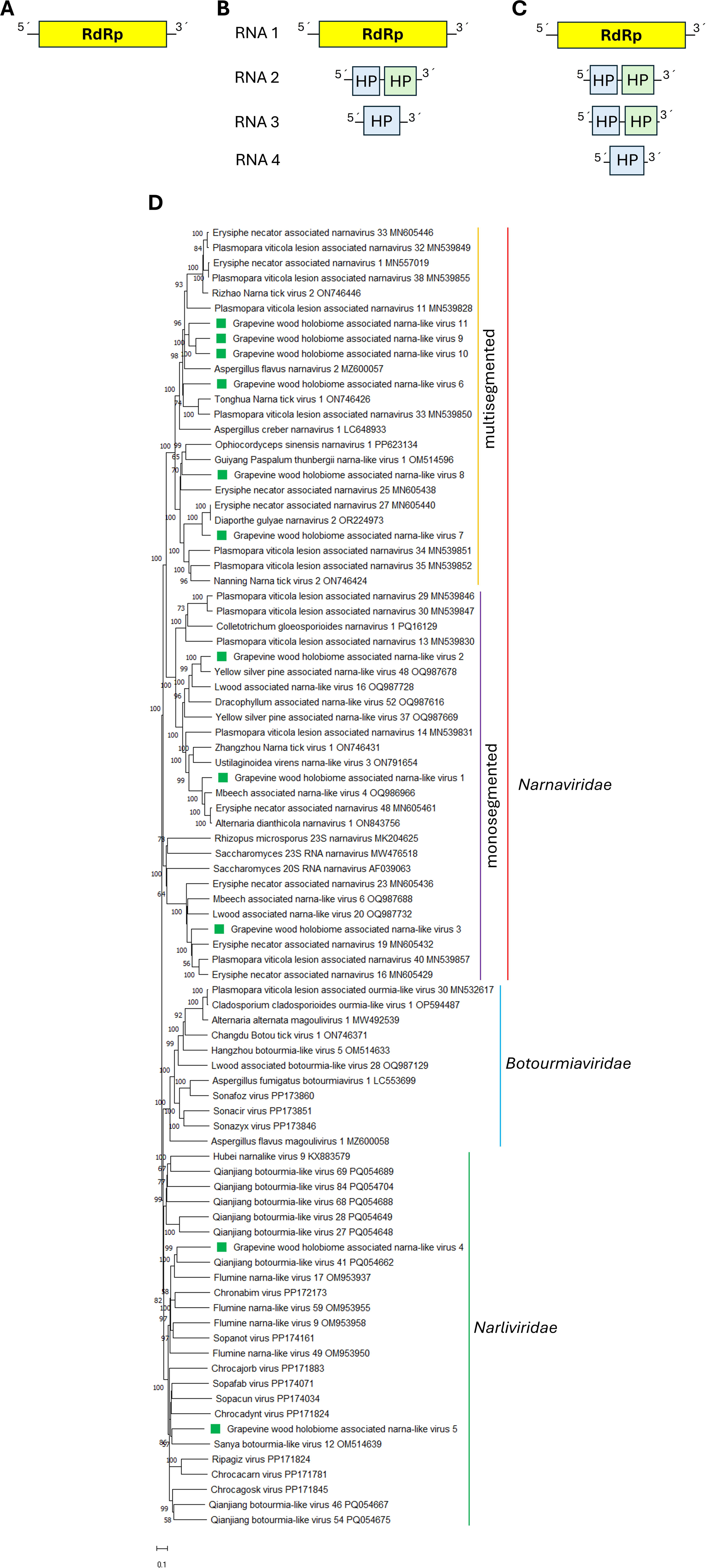
**(A)** Schematic representation of the genome organization of grapevine wood holobiome associated narna-like virus 1-5. **(B)** Schematic representation of the genome organization of grapevine wood holobiome associated narna-like virus 7-11. **(C)** Schematic representation of the genome organization of grapevine wood holobiome associated narna-like virus 6. **(D)** Maximum likelihood phylogenetic trees reconstructed using the RdRp protein sequence of grapevine wood holobiome associated narna-like virus 1-11 and of representative narnavirids. Botourmiavirus sequences were used as outgroup. Bootstrap values above 50% are shown (1000 replicates). The grapevine wood holobiome associated ambiguivirus narna-like virus 1-11 are indicated with a green square. The scale bar shows the substitution per site.

BLASTp searches using the deduced aa sequences showed that the best hits for GWHaNLV1-5 were Mbeech associated narna-like virus 4, Yellow silver pine associated narna-like virus 48, Erysiphe necator associated narnavirus 16, Qianjiang botourmia-like virus 41 and Sopafab virus RdRp’s respectively, with identity values below 82% (Table 1). Phylogenetic analysis based on RdRp aa sequence alignment revealed that GWHaNLV1-3 grouped with narna-like viruses likely belonging to the *Narnaviridae* family (Fig. 6D). Two major clades were observed within the monocistronic narnavirids, with GWHaNLV1 and GWHaNLV2 belonging to one clade, where GWHaNLV1 clustered with Mbeech associated narna-like virus 4, Alternaria dianthicola narnavirus 1 and Erysiphe necator associated narnavirus 48 (Fig. 6D), while GWHaNLV2 clustered with Yellow silver pine associated narna-like virus 48 (Fig. 6D). GWHaNLV3 belongs to the other clade (Fig. 6D) and grouped with Erysiphe necator associated narnavirus 16, Erysiphe necator associated narnavirus 19 and Plasmopara viticola lesion associated narnavirus 40 (Fig. 6D). Interestingly, the motifs MGD(MM/LF/LM) and GDD, previously described for Chiapello et al., (2020) as characteristic of a specific narnavirus group (e.g., Plasmopara viticola lesion associated narnavirus 13, 29 and 30), were also found in the RdRp of GWHaNLV1 and GWHaNLV2. In contrast, the motifs MGLG/P and GDD were present in the RdRp of GWHaNLV3 and other monosegmented narnaviruses belonging to the second major clade. Therefore, based on phylogenetic relationships and the presence of the MGD(MM/LF/LM) or MGLG/P motif, two genera could be proposed within the *Narnaviridae* family to accommodate monosegmented narnavirids. However, the demarcation criteria based on RdRp identity still need to be established. On the other hand, GWHaNLV4 and GWHaNLV5 grouped with narna-like viruses belonging to the clade of the proposed family “*Narliviridae*” (Sadiq et al., 2022) (Fig. 6D). GWHaNLV4 clustered with Qianjiang botourmia-like virus 41 within a clade of monocistronic narliviruses, while GWHaNLV5 clustered with Sanya botourmia-like virus 12 in another monocistronic narlivirus clade (Fig. 6D). Therefore, GWHaNLV4 and GWHaNLV5 could be classified within the same genus that accommodates monocistronic viruses in the proposed family “*Narliviridae*”. However, the demarcation criteria based on RdRp identity need to be established.

###### Multisegmented

Nineteen contigs showed similarity to two recently described multisegmented narna-like viruses. These multisegmented viruses were identified in *Aspergillus creber*, and four RNA segments were found (Chiba et al., 2021A). RNA 1 encodes the RdRp, while the other segments encode proteins of hypothetical function (Chiba et al., 2021A). The six identified multisegmented narna-like viruses were named as grapevine wood holobiome associated narna-like virus 6-11 (GWHaNLV6-11). The full-length coding sequence of four segments were assembled for GWHaNLV6 (Fig. 6C), and deposited in GenBank under accession numbers PV021971, PV021972, PV021973 and PV021974 (Table 1), while the full-length coding sequence of three segments were assembled for GWHaNLV7-11 (Fig. 6B) and deposited in GenBank under accession numbers PV021975-PV021989 (Table 1).

RNA 1 of the six viruses encodes an RdRp (Fig. 6B and 6C), and the aa identities among these six viruses ranged between 34.5.% to 74.2%. BLASTp searches of the RdRp showed that the best hits for GWHaNLV6-11 were Plasmopara viticola lesion associated narnavirus 32; Diaphorte gulyae narnavirus 2, Ophiocordyceps sinensis narnavirus 1, Erysiphe necator associated narnavirus 1, Erysiphe necator associated narnavirus 1 and Erysiphe necator associated narnavirus 1 (Table 1). RNA 2 of the six viruses encodes two hypothetical proteins (HP1 and HP2) (Fig. 6B and 6C), while RNA 3 of GWHaNLV6 encodes two HPs (Fig. 6C), whereas RNA 3 of GWHaNLV7-11 encode a single HP (Fig. 6B). RNA 4 of GWHaNLV6 encodes one HP (Fig. 6C) BLASTp searches of those HPs showed that the best hit was Aspergillus creber narnavirus 1 (AcreNV1) HPs (Table 1).

Phylogenetic analysis based on RdRp aa sequence alignment revealed that GWHaNLV6-11 formed a clade distinct from the monosegmented narnaviriuses clade (Fig. 6D). Two major clusters were observed within this clade: GWHaNLV6, GWHaNLV9-11 belonged to one mayor cluster, where GWHaNLV6 grouped with Plasmopara viticola lesion associated narnavirus 33 and Tonghua Narna tick virus (Fig. 6D), while GWHaNLV9-11 formed a subcluster with Aspergillus flavus narnavirus 2 (Fig. 6D). On other hand, GWHaNLV7-8 formed the other major cluster, where GWHaNLV7 grouped with Diaphorte gulyae narnavirus 2 and Erysiphe necator associated narnavirus 27 (Fig. 6D), while GWHaNLV8 grouped with Ophiocordyceps sinensis narnavirus 1 and Guiyang Paspalum thunbergii narna-like virus 1 (Fig 6D).

The RdRp of AcreNV1 and related plasmopara viticola lesion associated narnaviruses contain the MGDP motif (Chiapelo et al., 2020; Chiba et al., 2021), rather than the MGD(MM/LF/LM) or MGLG/P motifs found in the RdRp of monosegmented narnaviruses. The RdRp of GWHaNLV6-11 also contains the MGDP motif, further supporting their multisegmented genomic organization. Interestingly, only a single segment has been described for plasmopara viticola lesion associated narnaviruses and other viruses phylogenetically related to AcreNV1 and GWHaNLV6-11. Therefore, further analysis of the raw sequencing data should be conducted to identify potential missing segments of these viruses. A new viral family distinct from *Narnaviridae* could be proposed to accommodate these multisegmented narna-like viruses. The number of genera, likely two, and the demarcation criteria based on RdRp identity still need to be established.

##### Spliplamiviruses

Thirty-three contigs showed similarity to splipalmiviruses. These viruses were recently classified and accommodated within a new family *Splipalmiviridae*, including three genera (*Jakapalmivirus*, *Divipalmivirus*, and *Delepalmivirus*) (https://ictv.global/files/proposals/pending (<u>2024.003F.A.v1.Splipalmiviridae_newfam.docx</u>) (https://ictv.global/taxonomy) . Splipalmiviruses are phylogenetically related to narnavirids but have multi-segmented (+) RNA genomes that encode divided RdRps in two independent genomic segments (Chiba et al., 2021A-B; Sato et al., 2022; Kondo et al., 2022; Daghino et al., 2024). In these viruses, the MGDP motif is present in one RdRp segment, while the GDD motif is present in the other RdRp segment (Chiba et al., 2021A-B). The genome segments of the splipalmiviruses that do not encode the RdRp, i.e. RNA3 and RNA4, can be either mono- or polycistronic (Daghino et al., 2024). Moreover, splipalmiviruses assigned to the same tentative clade based on alignment of their RNA1- and RNA2-encoded proteins also exhibit the highest similarity in their RNA3-encoded proteins, with a few exceptions (Daghino et al., 2024).

The full-length coding sequence of three segments were assembled for GWHaNLV12-22 (Fig. 7A and B) and deposited in GenBank under accession numbers PV021990-PV022025 (Table 1). The RNA 1 and RNA2 segments of GWHaNLV12-22 encode the RdRp (Fig. 7A and B), while RNA 3 encodes a hypothetical protein. The RNA3 segment is polycistronic in GWHaNLV 12-16 and GWHaNLV18-19 (Fig. 7A) and monocistronic in GWHaNLV17 and GWHaNLV20-22 (Fig. 7B).

**Figure 7.**
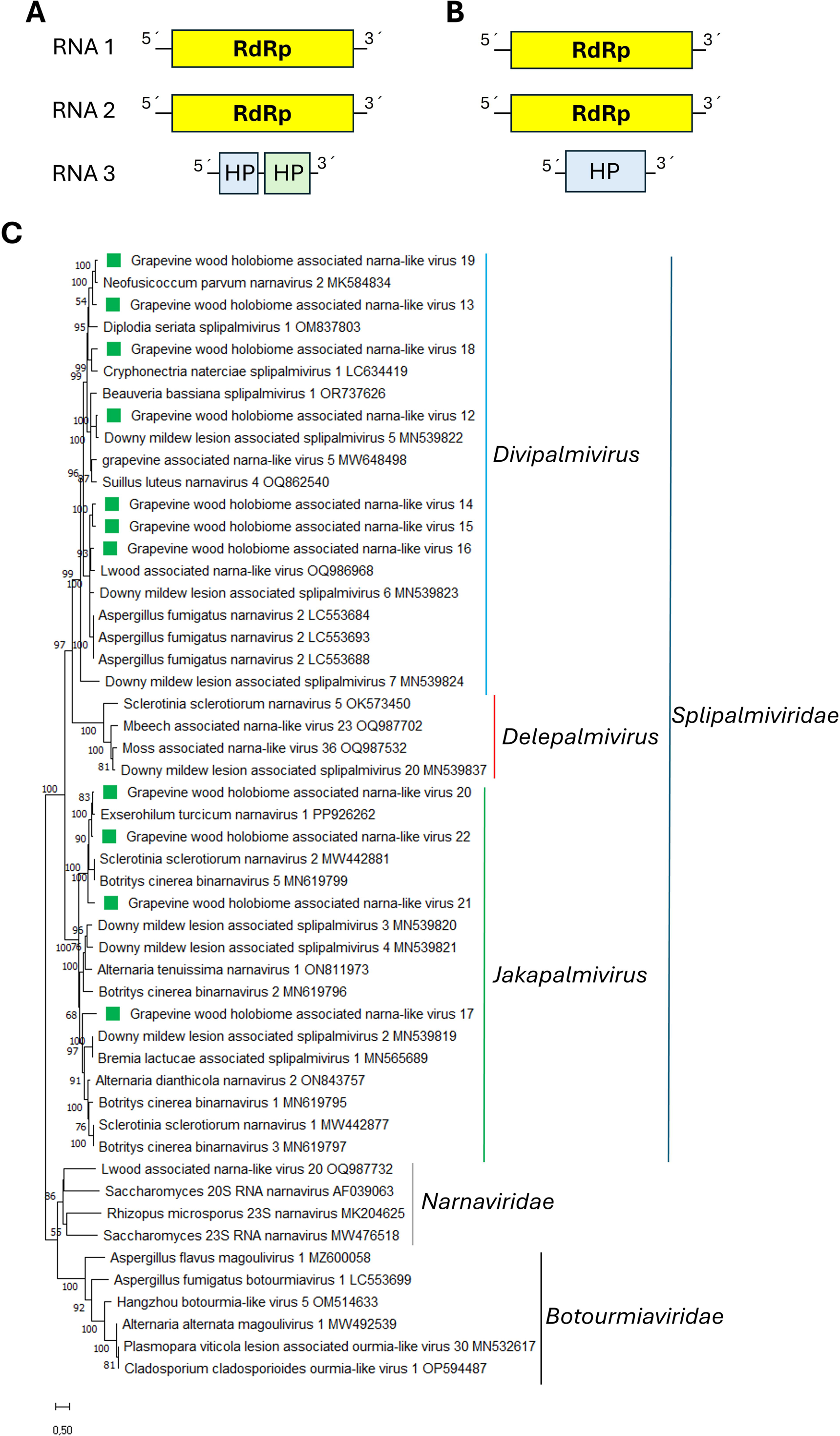
**(A)** Schematic representation of the genome organization of grapevine wood holobiome associated narna-like virus 12-16 and 18-19. **(B)** Schematic representation of the genome organization of grapevine wood holobiome associated narna-like virus 17 and 20-22. **(C)** Maximum likelihood phylogenetic trees reconstructed using the RdRp protein sequence of grapevine wood holobiome associated narna-like virus 11-22 and of representative splipalmiviruses. Narnavirus and Botourmiavirus sequences were used as outgroup. Bootstrap values above 50% are shown (1000 replicates). The grapevine wood holobiome associated narna-like virus 11-22 are indicated with a green square. The scale bar shows the substitution per site.

The aa identities of the RdRps encoded by segments 1 and 2 of the eleven viruses ranged from 23.3.% to 86.5%, while the highest aa identities between GWHaNLV12-22 and other spliplalmiviruses ranged from 47.9 to 89.39 (Table 1). BLASTp searches of the RdRp and HP showed that the best hits for GWHaNLV12-22 were their counterparts encoded by other splipalmiviruses (Table 1). As described by Daghino et al., (2024), the RNA 3-encoded proteins exhibited the highest similarity with those encoded by viruses belonging to the same clade.

Phylogenetic analysis based on RdRp aa sequence alignment revealed that GWHaNLV12-22 grouped into different clades within the family *Splipalmiviridae* (Fig. 7C). Seven viruses (GWHaOLV12-16 and GWHaOLV18-19) clustered with members of the genus *Divipalmivirus*, while the remaining four viruses (GWHaOLV17 and GWHaOLV20-22) clustered with members of the genus *Jakapalmivirus* (Fig 7C). Given the 90% aa RdRp identity threshold of demarcation criteria within the family all the splipalmiviruses reported here should be considered as members of new species within the *Divipalmivirus* and *Jakapalmivirus* genera.

##### Picorna-like viruses

Five contigs showed similarity to sequences from members of the order *Picornavirales,* which contain viruses with a monopartitite or bipartite positive-strand ssRNA genome that infect a wide range of hosts (Le Gall et al., 2008). This order comprises more than 323 species, classified into nine families and over 100 genera (Zell et al., 2022) (https://ictv.global/taxonomy). The five identified viruses were named grapevine wood holobiome associated picorna-like virus 1-5 (GWHaPLV1-5). The coding-complete sequences of GWHaPLV1, GWHaPLV2 and GWHaPLV4, as well as the partial sequences of GWHaPLV3 and GWHaPLV5, were obtained and deposited in GenBank under accession numbers PQ762193, PQ762194, PQ762196, PQ762195, and PQ762197, respectively (Table 1).

The coding-complete sequence of GWHaPLV1 is 9896nt in length and contains one ORF encoding a polyprotein of 3131aa (Fig. 8A). The Hel, Peptidase_C3, RdRp, rhv_like and CRPV_capsid domains were identified in its sequence. BLASTp searches of the nr protein database showed that the best hit was the polyprotein encoded by a Picornavirales sp. (coverage, 51%; E-value, 5e-154; identity, 43.07%). The coding-complete sequence of GWHaPLV2 is 9344 nt in length and has two ORFs (Fig. 8B). ORF1 encodes a putative CP of 823aa, containing the rhv_like and CRPV_capsid domains,while ORF2 encodes a putative non-structural polyprotein of 1956 aa, where the Hel and RdRp domains were identified. BLASTp searches showed that the best hit for both proteins were the CP encoded by a Riboviria sp (coverage, 98%; E-value, 2e-122; identity, 64,34%) and the RdRp protein encoded by Suncus murinus picorna-like virus 2 (coverage, 44%; E-value, 2e-73; identity, 44.79%), respectively. One ORF was predicted in the partially assembled sequence of GWHaPLV3 (Fig. 8A). BLASTp searches of the nr protein database showed that the best hit was the replicase encoded by a Riboviria sp. The coding-complete sequence of GWHaPLV4 is 9158 nt in length, and two ORFs were predicted (Fig. 8B). ORF1 encodes a putative CP of 879aa, where the domain rhv_like was identified, while ORF2 encodes a putative non-structural polyprotein of 1805aa, where the Hel and RdRp domains were identified. BLASTp searches showed that the best hit for both proteins were the CP encoded by a Riboviria sp (coverage, 97%; E-value, 4e-78; identity, 62.84%) and RdRp protein encoded by a Riboviria sp. (coverage, 49%; E-value, 1e-131; identity, 54.93%), respectively. Two ORFs were predicted in the partially assembled sequence of GWHaPLV5 (Fig. 8B). ORF1 encodes a putative CP of 825aa, containing the rhv_like and CRPV_capsid domains,while ORF2 encodes a putative non-structural polyprotein, which was partially assembled, where the Hel and RdRp domains were identified. BLASTp searches showed that the best hit for both proteins were the CP and RdRp protein encoded by a Dicistroviridae sp. (coverage, 98%; E-value, 0.0; identity, 51.34% for the CP). The Hel and RdRp domains form the replicative module, which is a typical feature of picornavirids (Zell et al., 2022). In GWHaPLV1 and GWHaPLV3, the CP is located in the C-terminal region, while in GWHaPLV2, GWHaPLV4 and GWHaPLV5 the CP is located in the N-terminal region of the virus. This arrangement is characteristic of picornavirids, where the CPs are either encoded together with the nonstructural proteins as part of a large polyprotein or by a second ORF (Zell et al., 2022).

**Figure 8.**
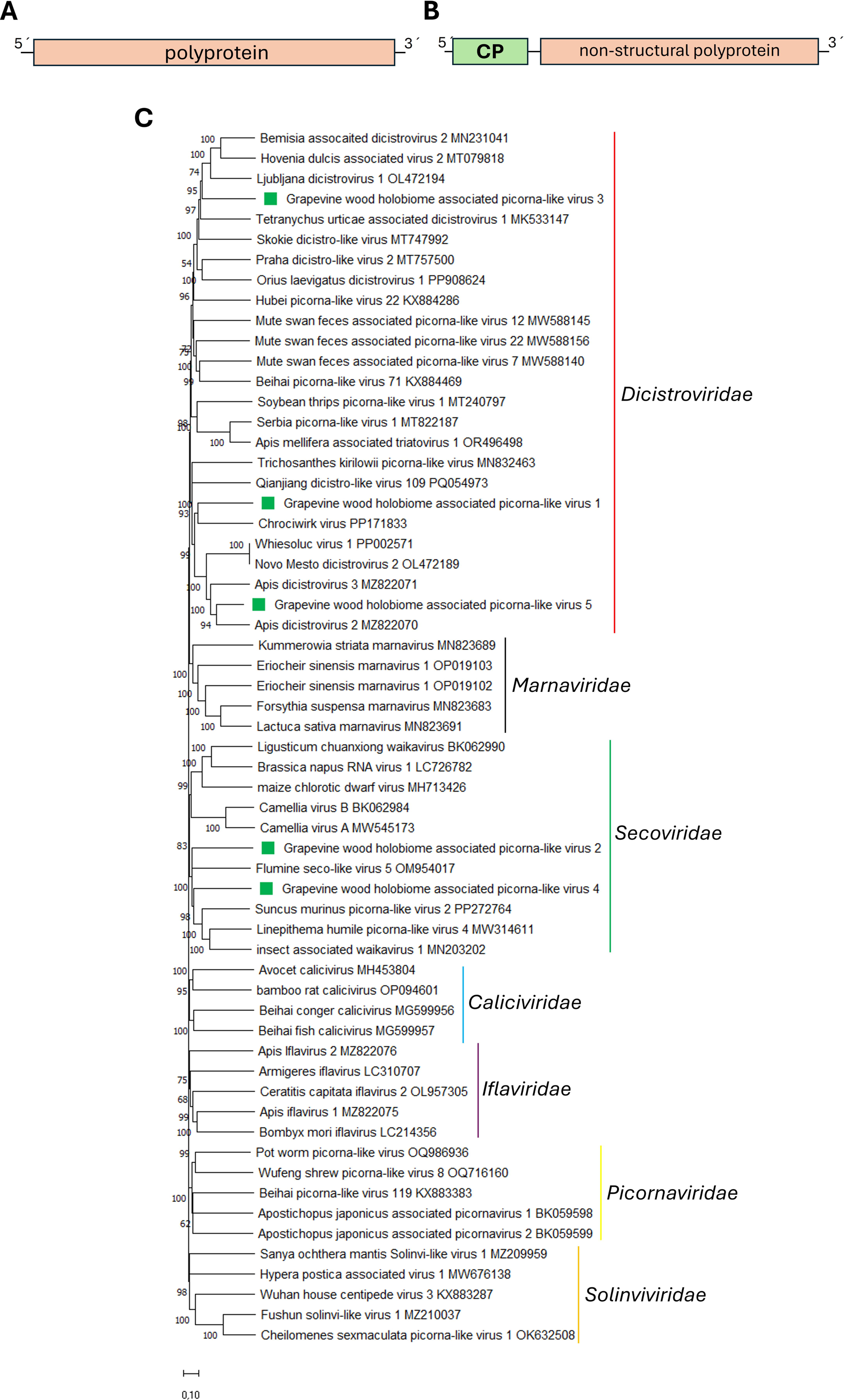
(**A**) Schematic representation of the genome organization of grapevine wood holobiome associated picorna-like virus 1 and 3 (**B**) Schematic representation of the genome organization of grapevine wood holobiome associated picorna-like virus 2 and 4-5. (**C**) Maximum likelihood phylogenetic trees reconstructed using the RdRp protein sequence of grapevine wood holobiome associated picorna-like virus 1-5 and of representative picornavirids. Bootstrap values above 50% are shown (1000 replicates). The grapevine wood holobiome associated picorna-like virus 1-5 are indicated with a green square. The scale bar shows the substitution per site.

Phylogenetic analysis based on RdRp aa sequence alignment revealed that GWHaPLV1, GWHaPLV2, GWHaPLV3, GWHaPLV4 and GWHaPLV5 formed a well-supported clade with other picornavirids (Fig. 8C). GWHaPLV1, GWHaPLV3 and GWHaPLV5 clustered with members of the *Dicistroviridae* family, while GWHaPLV2 and GWHaPLV4 clustered with members of the *Secoviridae* family (Fig. 8C). Therefore, GWHaPLV1, GWHaPLV2, GWHaPLV3, GWHaPLV4 and GWHaPLV5 are five novel putative picornavirids. GWHaPLV1 and GWHaPLV3 are monocistronic disctrovirids and should be classified into a genus of this family composed of monocistronic members, while GWHaPLV5 is a bi-cistronic dicistrovirid that should be accommodated into a genus of this family composed of bi-cistronic members. On the other hand, GWHaPLV2 and GWHaPLV4 are secovirids that belong to a clade of insect-associated secovirids (Fig 8C). It is tempting to speculate that GWHaPLV1, GWHaPLV2, GWHaPLV3, GWHaPLV4 and GWHaPLV5 are insect-associated viruses rather than fungal-associated ones, possibly originating from insect RNA co-purified with fungal samples.

##### Tombusvirus

One contig showed similarity to sequences from members of the family *Tombusviridae*, which includes viruses with non-segmented, positive-sense ssRNA genomes that typically contain 4-7 ORFs (Lozier et al., 2024). This family includes over 100 species classified into eighteen genera within three subfamilies (Lozier et al., 2024). The identified virus was named grapevine wood holobiome associated tombus-like virus (GWHaTLV). Its partial sequence was obtained and deposited in GenBank under accession number PV036960.

The partial coding-complete sequence of GWHaTLV is 4518 nt in length (Table 1), with five predicted ORFs (Fig 9A), where the first ORF is incompletely assembled, lacking coverage at its 5’ region. The genomic organization of GWHaTLV is similar to that of Erysiphe necator associated tombus-like virus 2 (MN627442), Nanning tombu tick virus 1 (ON746539), among other tombusvirids (Lozier et al., 2024). The presumed ORF1 is immediately followed by ORF2, which encodes the putative RdRp. The best hit in BLASTp searches of the nr protein database was the Erysiphe necator associated tombus-like virus 2 RdRp (coverage, 100%; E-value, 0.0; identity, 86.65%). ORF2 is translated via in-frame readthrough of the stop codon of the preceding ORF, beginning immediately after the stop codon rather than at the first AUG codon of the ORF. This arrangement is characteristic of tombusvirids (Lozier et al., 2024). ORF3 encodes a putative CP with a size of 214 aa, whose best hit in BLASTp searches was the Zizania latifolia tombusvirus CP (coverage, 99%; E-value, 1e-140; identity, 90.1%). ORF5 is located within the same reading frame and appears to be translated via readthrough of the ORF3 stop codon, generating a 376-aa protein that serves as a terminal extension of the CP. The organization of ORFs 3, 4 and 5 is similar to that reported for maize-associated tombusvirus, and related viruses, which have been proposed for classification within the genus “*Rimosavirus*”. This organization also resembles that of reported for luteoviruses and poleroviruses (Lozier et al., 2024). The ORF4, which overlaps with ORF3, encodes a putative HP of 236 aa, whose best hit in the BLASTp search was the Nanning tombu tick virus 1 HP (coverage, 78%; E-value, 3e-12; identity, 33.5%).

Phylogenetic analysis based on RdRp aa sequence alignment revealed that GWHaTLV clustered within a well-supported clade of tombus-like viruses proposed to be assigned within the “*Rimosavirus*” genus, which includes plant-, fungal-, invertebrate- and vertebrate-associated viruses (Lozier et al., 2024) (Fig. 9B). Specifically, GWHaTLV grouped with Erysiphe necator associated tombus-like virus 2, Chrocowerd virus and Chrocotusk virus, (Fig. 9B). Therefore, GWHaTLV is a novel member of the *Tombusviridae* family and could be classified within the “*Rimosavirus*” genus. However, demarcation criteria based on the identity of the RdRp and/or CP proteins need to be established for this genus. Rimosaviruses have a wide host range, but it has been hypothesized that they are plant pathogens (Lozier et al., 2024). Thus, it is likely that GWHaTLV1 is a grapevine-associated virus rather than a mycovirus.

**Figure 9.**
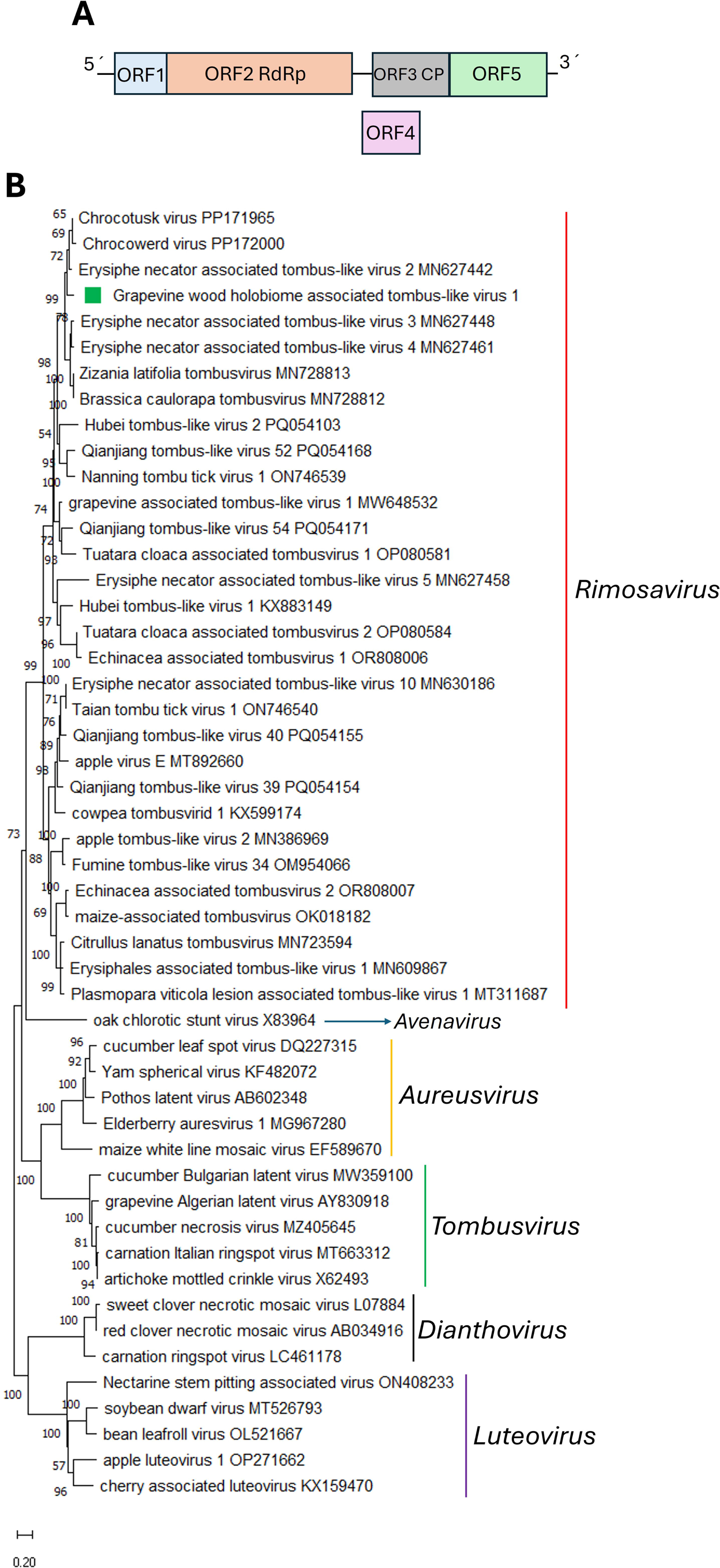
(**A**) Schematic representation of the genome organization of grapevine wood holobiome associated tombus-like virus. (**B**) Maximum likelihood phylogenetic trees reconstructed using the RdRp protein sequence of grapevine wood holobiome associated tombus-like virus and of representative tombusvirids. Bootstrap values above 50% are shown (1000 replicates). The grapevine wood holobiome associated tombus-like virus is indicated with a green square. The scale bar shows the substitution per site.

### 3.2B. Negative-sense single-stranded RNA viruses

#### Bunya-like viruses

Three contigs showed similarity to sequences from members of the order *Hareavirales*, class *Bunyaviricetes,* which includes viruses that infect a wide range of hosts, including human, mammals, invertebrates, and plants. Most of these viruses share a similar tri-segmented, negative-sense ssRNA genomic organization (Leventhal et al., 2021; Kuhn et al., 2024). The two identified bunyavirids were named as grapevine wood holobiome associated bunya-like virus 1-2 (GWHaBLV1-2). Two segments were obtained for GWHaBLV1 and deposited in GenBank under accession numbers PQ683342 and PQ683343, while for GWHaBLV2, only one segment was assembled and deposited under accession number PQ683344.

For GWHaBLV1, RNA1 was nearly completely assembled, with a sequence length of 6440 nt, while the complete-coding region of RNA2 was fully assembled, consisting of 1016 nt. On the other hand, for GWHaBLV2, only RNA 1 was partially assembled (Table 1).

RNA1 of both viruses contains a single ORF encoding an RdRp (Fig. 10A). BLASTp searches of the nr protein database using the GWHaBLV1 RdRp showed that the best hits were Sclerotinia sclerotiorum bunyavirus 5, Sclerotinia sclerotiorum bunyavirus 6 and Sclerotinia sclerotiorum bunyavirus 7 (SsBYB5-7), with identities values ranging from 63.47 to 64.68%. These viruses have been proposed for classification within the family “Mycophenuiviridae” (Jia et al., 2021). Moreover, the eight conserved motifs that represent highly conserved central regions of RdRps from members of the order Bunyavirales and were found in the SsBYB5-7 RdRp’s (Jia et al., 2021), were also identified in the GWHaBLV1 RdRp. Additionally, the conserved endonuclease domain and motif G, previously identified in the SsBYB5-7 RdRps (Jia et al., 2021), were also present in the GWHaBLV1 RdRp. On the other hand, BLASTp searches using the GWHaBLV2 RdRp showed that the best hit was the bunyavirid Plasmopara viticola lesion associated mycobunyavirales-like virus 3 RdRp. RNA 2 of GWHaBLV1 contains a single ORF encoding a nucleocapsid protein (NC) of 211aa (Fig. 10A), in which the Tenui_N conserved domain was identified. Interestingly, BLASTp searches showed that the best hit for this NC was the plant-associated Tulip streak virus NC (coverage, 89%; E-value, 4e-25; identity, 34.25%). Only one segment has been identified in SsBYB5-7 (Jia et al., 2021), thus, it is likely that additional segments of these viruses could be identified if sequencing data is reanalyzed.

**Figure 10.**
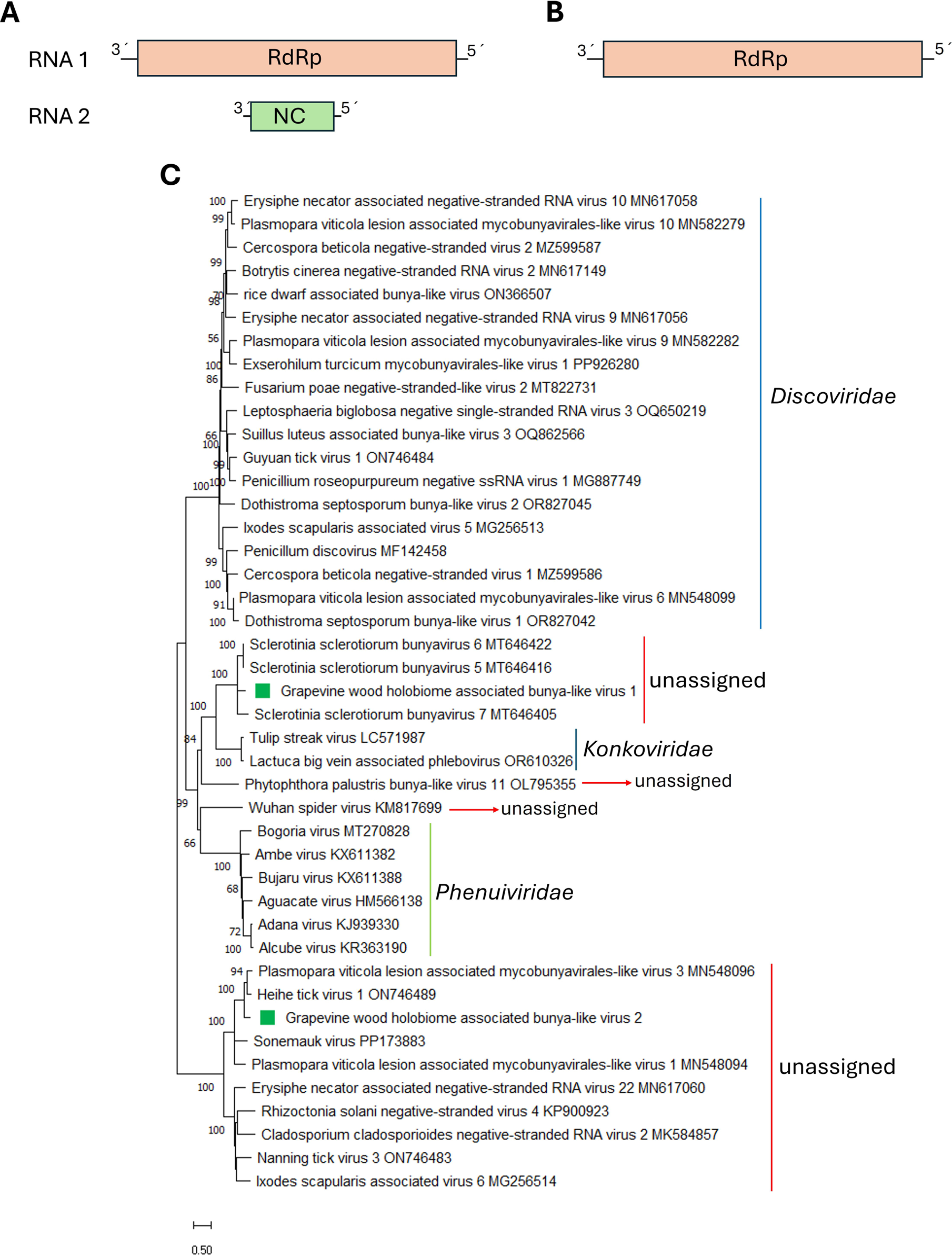
(**A**) Schematic representation of the genome organization of grapevine wood holobiome associated bunya-like virus 1. (**B**) Schematic representation of the genome organization of grapevine wood holobiome associated bunya-like virus 2. (**C**) Maximum likelihood phylogenetic trees reconstructed using the RdRp protein sequence of grapevine wood holobiome associated bunya-like virus 1-2 and of representative bunyavirids. Bootstrap values above 50% are shown (1000 replicates). The grapevine wood holobiome associated bunya-like virus 1-2 are indicated with a green square. The scale bar shows the substitution per site.

The phylogenetic tree based on RdRp aa sequence alignment placed GWHaBLV1 and GWHaBLV2 with other bunyavirids in different clusters (Fig. 10B). GWHaBLV1 clustered with SsBYB5-7, forming a clade related to the unassigned Phytopthora palustris bunya-like virus 11, which also has two RNA segments (Botella et al., 2022). This clade is further related to the cluster containing members of the family *Konkoviridae*, which was recently created to accommodate plant-associated viruses (Neriya and Nishigawa, 2024). Members of *Konkoviridae* have four RNA segments, where the third segment encodes a putative movement protein, while the fourth encodes a protein of unknown function. No additional segments were identified in GWHaBLV1, suggesting that these two additional segments are not essential for its genome and that this virus is exclusively fungi-associated. Supporting this assumption, recent studies have identified phenuiviruses related to plant-associated members of the genus *Laulavirus*, that were isolated from fungi, with three segments, including one encoding a putative movement protein. These viruses can infect both plant and fungal hosts, redefining the conventional distinction between fungal and plant viruses (Dai et al., 2024). On the other hand, GWHaBLV2 clustered separately with the unassigned viruses Plasmopara viticola lesion associated mycobunyavirales-like virus 3 and Heihe tick virus 1 (Fig. 10B). Interestingly, all viruses in this cluster, including GWHaBLV2, have only one identified segment. Thus, it is highly likely that the genome descriptions of GWHaBLV2 and related viruses are incomplete. Both GWHaBLVV1 and GWHaBLV2 represent novel bunyavirid members that should be assigned to two newly proposed families to be created within the order *Hareavirales*, class *Bunyaviricetes*.

#### Discoviruses

Five contigs showed similarity to sequences from members of the recently established family *Discoviridae*, order *Hareavirales*, which comprises negative-sense ssRNA fungi-associated viruses with genomes consisting of three monocistronic RNA segments (Kuhn et al., 2023). The five identified segments correspond to two putative viruses, which were named as grapevine wood holobiome associated discovirus 1-2 (GWHaDV1-2). For GWHaDV1, all three segments were obtained and deposited in GenBank under accession numbers PQ508371, PQ508372 and PQ508373 while for GWHaDV2, only two segments were assembled (RNA 2 and RNA3) and deposited in GenBank under accession numbers PQ508374 and PQ508375, leaving its genome incomplete (Table 1).

The RNA1, RNA2, and RNA3 segments of GWHaDV1 have 6502, 1821, and 1195 nt in length, respectively (Fig. 11A), while the RNA 2 and RNA3 segments of GWHaDV2 have 1563 and 1135 nt in length (Fig. 11B). RNA1 of GWHaDV1 encodes the large protein (L), which is 2138 aa in lenght, and contains the RDRP_SSRNA_NEG_SEG conserved domain. BLASTp searches of the nr protein database showed that the best hit was the L protein encoded by Leptosphaeria biglobosa negative single-stranded RNA virus 3 (coverage, 99%; E-value, 0.0; identity, 65.19%). RNA 2 of both GWHaDV1 and GWHaDV2 encodes a non-structural protein (NS) of 557 aa and 482 aa, respectively. BLASTp searches showed that the best hits were the NS encoded by Leptosphaeria biglobosa negative single-stranded RNA virus 3 (coverage, 94%; E-value, 2e-92; identity, 33.70%) and Plasmopara viticola lesion associated mycobunyavirales-like virus 4 (coverage, 95%; E-value, 2e-54; identity, 30.68%), respectively. RNA 3 of both GWHaDV1 and GWHaDV2 encodes a nucleocapsid protein (NC) of 305 aa and 285 aa, respectively. BLASTp searches showed that the best hits were the NC encoded by Leptosphaeria biglobosa negative single-stranded RNA virus 9 (coverage, 90%; E-value, 4e-133; identity, 66.55%), and Plasmopara viticola lesion associated mycobunyavirales-like virus 4 (coverage, 100%; E-value, 2e-175; identity, 79.65%), respectively (Table 1).

**Figure 11.**
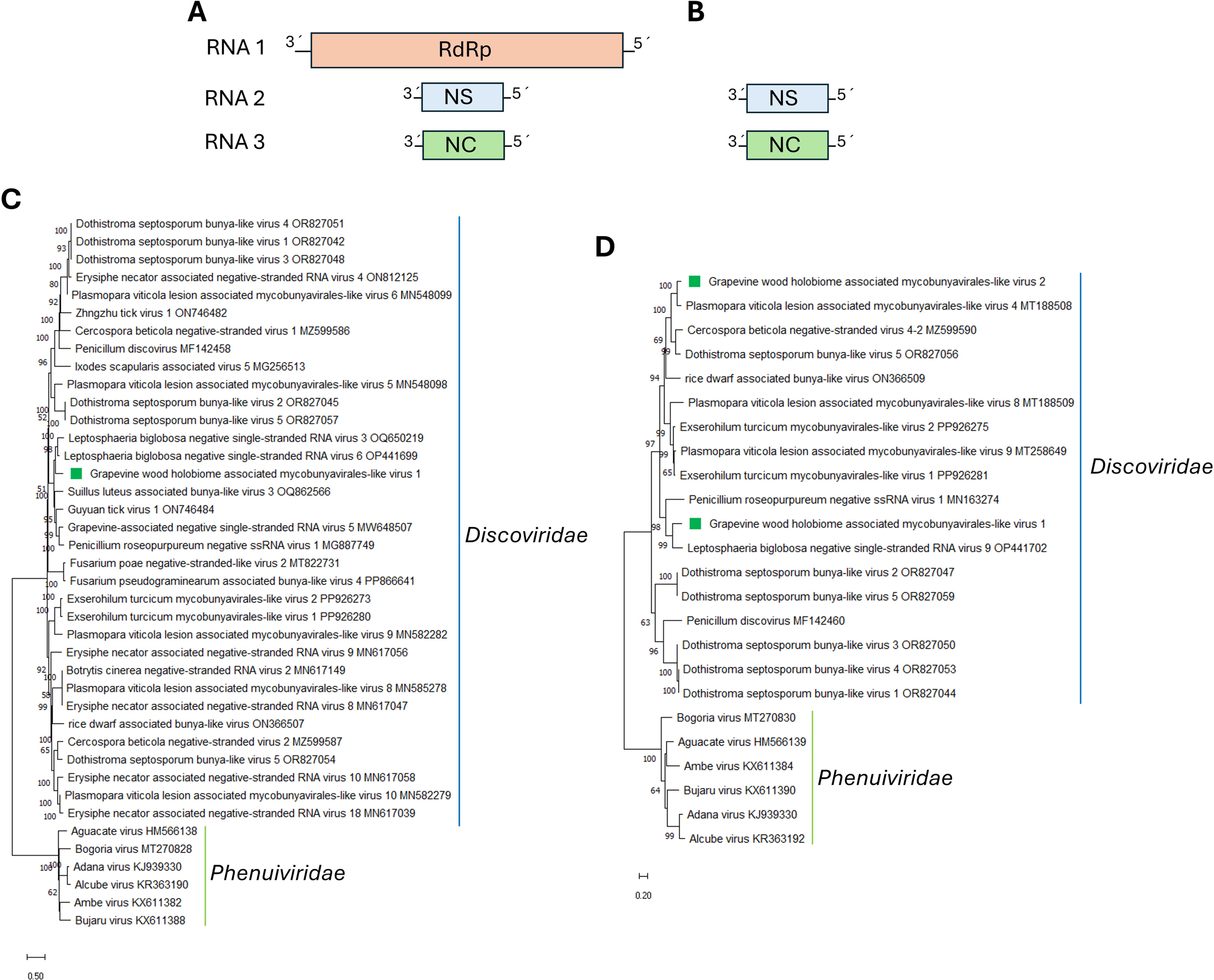
(**A**) Schematic representation of the genome organization of grapevine wood holobiome associated discovirus 1. (**B**) Schematic representation of the genome organization of grapevine wood holobiome associated discovirus 2. (**C**) Maximum likelihood phylogenetic trees reconstructed using the RdRp protein sequence of grapevine wood holobiome associated discovirus 1 and of representative discoviruses. (**D**) Maximum likelihood phylogenetic trees reconstructed using the NC protein sequence of grapevine wood holobiome associated discovirus 1-2 and of representative discoviruses. Phenuivirids sequences were used as outgroup. Bootstrap values above 50% are shown (1000 replicates). The grapevine wood holobiome associated discovirus 1-2 are indicated with a green square. The scale bar shows the substitution per site.

The phylogenetic tree based on RdRp aa sequence alignment placed GWHaDV1 within a cluster of discoviruses, alongside Leptosphaeria biglobosa negative single-stranded RNA virus 3 and Leptosphaeria biglobosa negative single-stranded RNA virus 6 (Fig. 11C). Meanwhile, the phylogenetic tree based on NC aa sequence alignment placed GWHaDV1 and GWHaDV2 in different clusters within the *Discoviridae* family (Fig. 11D). GWHaDV1 grouped with Leptosphaeria biglobosa negative single-stranded RNA virus 9 and Penicillum roseopurpureum negative ssRNA virus 1, while GWHaDV2 clustered separately with Plasmopara viticola lesion associated mycobunyavirales-like virus 4 (Fig. 11D). According to the ICTV species demarcation criteria, classification should be based on sequence identity in the complete RdRp, with a threshold of less than 80% amino acid sequence identity (Kuhn et al., 2023). Therefore, GWHaDV1 represents a novel member of the *Orthodiscovirus* genus within the *Discoviridae* family.

#### Mononegaambiviruses

Four contigs showed similarity to sequences from members of the recently created family *Mymonaviridae*, order *Mononegavirales*, which comprises viruses with negative-sense ssRNA genomes (Jiang et al 2022). The family includes nine genera, where viruses from eight genera have monopartite genomes, while those belonging to the genus *Penicillimonavirus* are bipartite (Moran et al., 2023). The two identified mymonaviruses were named as grapevine wood holobiome associated mononegaambi virus 1-2 (GWHaMV1-2). The coding-complete sequence of the two segments of both viruses were obtained and deposited in GenBank under accession numbers PQ643870 and PQ643871 and PQ643872 and PQ643873, respectively (Table 1).

Both viruses have two RNA segments (Fig. 12A), and their organization is similar to that reported for the penicillimonaviruses Plasmopara viticola lesion associated mononegaambi virus 2, Plasmopara viticola lesion associated mononegaambi virus 4 (Chiapello et al., 2020), Plasmopara viticola lesion associated mononegaambi virus 3 (Moran et al., 2023) and Exserohilum turcicum mymonavirus 1 (PP926276), among others. The coding-complete sequence of GWHaMV1 RNA1 is 6700 nt in length, while that of GWHaMV2 RNA1 is 6777 nt, and hey share 68.7% identity. Both viruses showed the highest sequence identity of 60.9% and 61.4%, respectively, with Plasmopara viticola lesion associated mononegaambi virus 2. The ORF1 of GWHaMV1 and GWHaMV2, located on the positive-sense strand, encodes a nucleocapsid protein (NC) (Fig. 12A) of 189 aa and 186 aa, respectively, sharing a 48.1% identity. The results of BLASTp searches of the nr protein database showed that the best hit was the NC encoded by Plasmopara viticola lesion associated mononegaambi virus 2 (coverage, 79%; E-value, 5e-25; identity, 38.67%, and coverage, 82%; E-value, 9e-21; identity, 34.64%, respectively). The ORF2 of GWHaMV1 and GWHaMV2, located on the negative-sense strand, encodes an RdRp (Fig. 12A) of 1944 aa, sharing 84.3% identity. BLASTp searches showed that the best hit was the RdRp encoded by Plasmopara viticola lesion associated mononegaambi virus 2 (coverage, 99%; E-value, 0.0; identity, 69.65%, and coverage, 99%; E-value, 0.0; identity, 69.26%, respectively) (Table 1). The conserved domains Mononeg_RNA_pol and Mononeg_mRNAcap were identified in the RdRps of GWHaMV1 and GWHaMV2.

**Figure 12.**
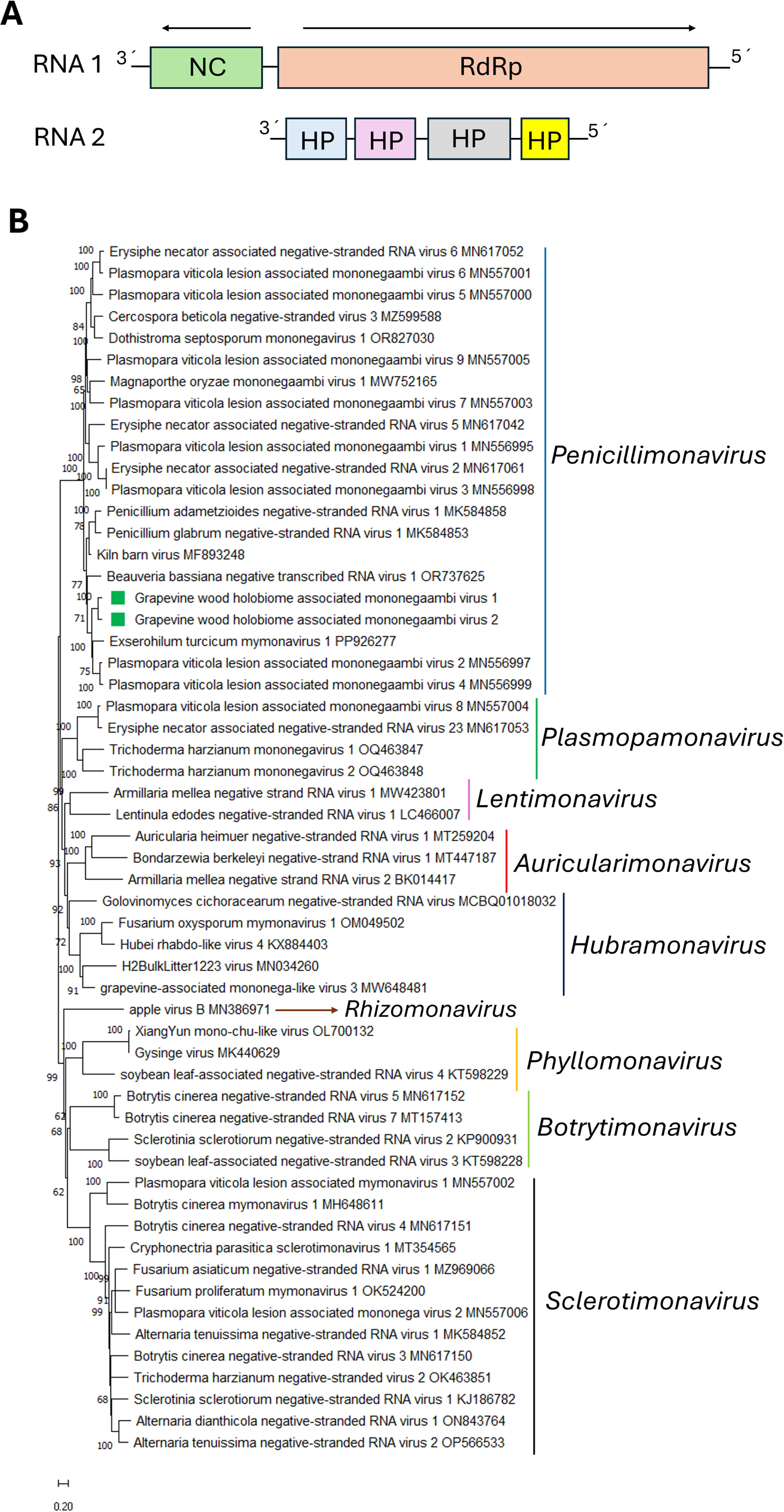
(**A**) Schematic representation of the genome organization of grapevine wood holobiome associated mononegaambi virus 1-2. (**B**) Maximum likelihood phylogenetic trees reconstructed using the RdRp protein sequence of grapevine wood holobiome associated mononegaambi virus 1-2 and of representative mymonaviruses. Bootstrap values above 50% are shown (1000 replicates). The grapevine wood holobiome associated mononegaambi virus 1-2 are indicated with a green square. The scale bar shows the substitution per site.

The coding-complete sequence of GWHaMV1 RNA2 is 3648 nt in length, while that of GWHaMV2 RNA2 is 3664 nt (Table 1). Both GWHaMV1 and GWHaMV2 RNA2 contain four open reading frames (ORF1, ORF2, ORF3 and ORF4) (Fig. 12A). ORF1 encodes an hypothetical protein (HP) of 260 and 259 aa, respectively, which the best hit being the HP encoded by Exserohilum turcicum mymonavirus 1. ORF2 encodes an HP of 391 aa, which the best hit being the HP encoded by Plasmopara viticola lesion associated mononegaambi virus 2. ORF3 encodes an HP of 205 aa but did not return any significant hits in NCBI database. Similarly, ORF 4 encodes an HP of 165, also with no hits in the NCBI database.

The phylogenetic tree based on RdRp aa sequence alignment placed GWHaMV1 and GWHaMV2 within a well-supported clade of penicillimonaviruses (Fig. 12B). Both viruses clustered with Beuaveria bassiana negative transcribed RNA virus 1, Plasmopara viticola lesion associated mononegaambi virus 2, Plasmopara viticola lesion associated mononegaambi virus 4, and Exserohilum turcicum mymonavirus 1 (Fig. 12B). According to ICTV classification criteria, members of different species within the genus *Penicillimonavirus* must differ by more than 30% in nucleoprotein amino acid sequence and by more than 30% in their coding-complete or complete genome nucleotide sequence (Jiang et al., 2022). Therefore, both GWHaMV1 and GWHaMV2 represent novel members of the genus *Penicillimonavirus* within the family *Mymonaviridae*.

#### Mycoophioviruses

Four contigs showed similarity to sequences from members of the recently proposed *Myco*a*spiviridae* family, order *Nadrevirales*, which consist of viruses with negative-sense ssRNA genomes that are fungi-associated (Chiapello et al., 2020). In contrast to their plant-associated counterparts, most of the identified mycoophiovirus to date have been found to possess a single genome segment (Chiapello et al., 2020; Liu et al., 2023B). However, a second genome segment has been identified in mycoophioviruses associated to Colletotrichum (Hamim et al., 2022) and Fusarium (Buivydaite et al., 2024). Therefore, it is tempting to speculate that all mycoophioviruses may possess more than one genomic segment, similar to plant-associated ophioviruses (Liu et al., 2023B). The two identified mycoophioviruses were named as grapevine wood holobiome associated mycoophiovirus 1-2 (GWHaMOV1-2). The coding-complete sequence of both segments for each virus were obtained and deposited in GenBank under accession numbers PQ638356 and PQ638357 for GWHaMOV1, PQ638358 and PQ638359 for GWHaMOV2 (Table 1).

Both GWHaMOV1 and GWHaMOV2 genomes consist of two segments (Fig. 13A), similarly to those mycoophioviruses associated to Colletotrichum (Hamim et al., 2022) and Fusarium (Buivydaite et al., 2024), which is consistent with the suggestion by Chiapello et al., (2020) that mycoophioviruses might have additional genomic segments. The coding-complete sequence of the first segment of GWHaMOV1 is 7696 nt in length, while that of GWHaMOV 2 is 7651 nt. This segment contains a single open reading frame encoding a putative RdRp of 2359 aa (Fig. 13A), which are 69% identical. BLASTp searches of the nr protein database showed that the best hits for the RdRp of GWHaMOV1 and GWHaMOV2’s were the RdRp encoded by Leptosphaeria biglobosa negative single-stranded RNA virus 7 (coverage, 100%; E-value, 0.0; identity, 83.38%), and Cladosporium cladosporioides negative-stranded RNA virus 1 (coverage, 99%; E-value, 0.0; identity, 69.19%), respectively (Table 1). Moreover, the Mononeg_RNA_pol conserved domain was identified in the RdRps of GWHaMOV1 and GWHaMOV2, and all five conserved motifs shared by mycoophioviruses (Hamim et al., 2022; Liu et al., 2023B) were also identified in the RdRps of GWHaMOV1 and GWHaMOV2. The coding-complete sequence of the second segment of GWHaMOV1 is 2911 nt in length, while that of GWHaMOV 2 is 2872 nt. The second segment of GWHaMOV1 contains three OFRs encoding three hypothetical proteins (HP) of 108 aa, 400 aa and 328 aa (Fig. 13A). BLASTp searches of the nr protein database showed no hits for the first HP, while the best hit for HP2 was an HP encoded by Colletotrichum associated negative-sense single-stranded RNA virus 2 (coverage, 85%; E-value, 8e-102; identity, 47.55%), and the best hit for the HP3 was an HP encoded by Colletotrichum associated negative-sense single-stranded RNA virus 2 (coverage, 97%; E-value, 7e-169; identity, 69.69%) (Table 1). The second segment of GWHaMOV2 contains three OFRs encoding three HP of 108 aa, 393 aa and 328 aa (Fig. 13A). The results of a BLASTp search showed no hits for the first HP, while the best hit for HP2 was an HP coded by Colletotrichum associated negative-sense single-stranded RNA virus 2 (coverage, 83%; E-value, 7e-99; identity, 47.79%); and the best hit for the HP3 was an HP coded by Colletotrichum associated negative-sense single-stranded RNA virus 2 (coverage, 96%; E-value, 3e-167; identity, 69.81%) (Table 1). The HP3 encoded by both viruses showed identity to the protein encoded by tORF4 of the second RNA segment of Fusarium culmorum mycoophioviorus 1. Like that protein (Buivydaite et al., 2024), the HP3 encoded by GWHaMOV1 and GWHaMOV2 exhibited strong similarity to the nucleocapsid proteins of *Aspiviridae* members in HHpred searches. Thus, the HP3 encoded by GWHaMOV1 and GWHaMOV2 is likely a nucleocapsid protein.

**Figure 13.**
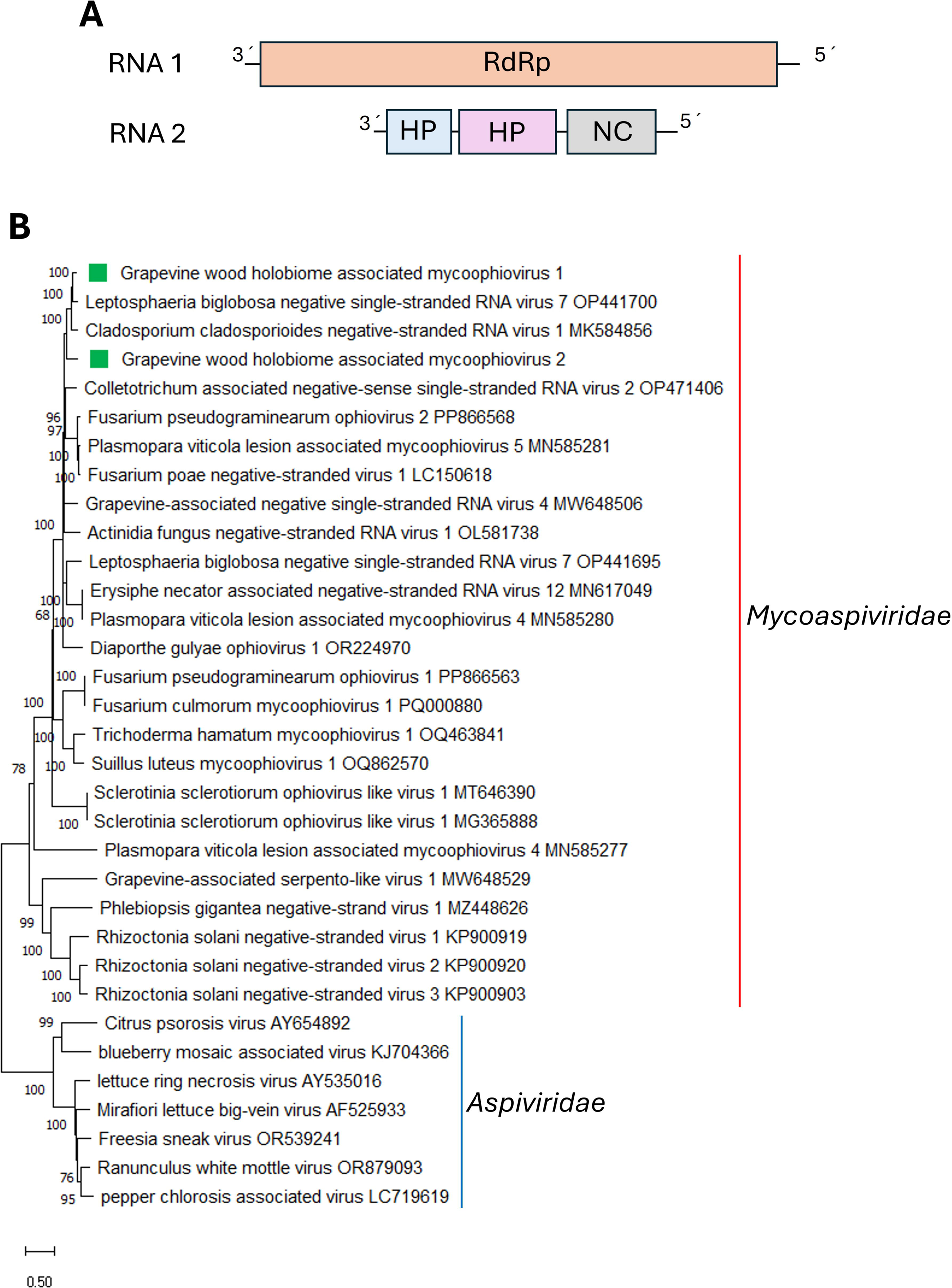
(**A**) Schematic representation of the genome organization of grapevine wood holobiome associated mycoophiovirus 1-2. (**B**) Maximum likelihood phylogenetic trees reconstructed using the RdRp protein sequence of grapevine wood holobiome associated mycoophiovirus 1-2 and of representative fungi and plant-associated aspivirids. The grapevine wood holobiome associated mycoophiovirus 1-2 are indicated with a green square. The scale bar shows the substitution per site.

A phylogenetic tree based on RdRp aa sequence alignment placed GWHaMOV1 and GWHaMOV2 within a well-supported clade of mycoaspivirids (Fig. 13B). Both viruses clustered with Leptosphaeria biglobosa negative single-stranded RNA virus 7 and Cladosporium cladosporioides negative-stranded RNA virus 1. Therefore, GWHaMOV1 and GWHaMOV2 represent novel members of the recently proposed family *Mycoaspiviridae* (Chiapello et al., 2020). The demarcation criteria based on RdRp protein identity still need to be established, along with the genera to be created within this proposed family.

#### Yue-like viruses

One contig showed similarity to yue-like virus sequences. The identified virus was named grapevine wood holobiome associated yue-like virus (GWHaYLV). The partial sequence of this virus was obtained and deposited in GenBank under the accession number PQ720401 (Table 1).

The partial sequence of GWHaYLV contains a single ORF encoding a putative RdRp (Fig. 14A). The results of a BLASTp search of the nr protein database showed that the best hit was the RdRp encoded by Goldenrod fern yue-like virus (coverage, 96%; E-value, 2E-60; identity, 29.04%).

**Figure 14.**
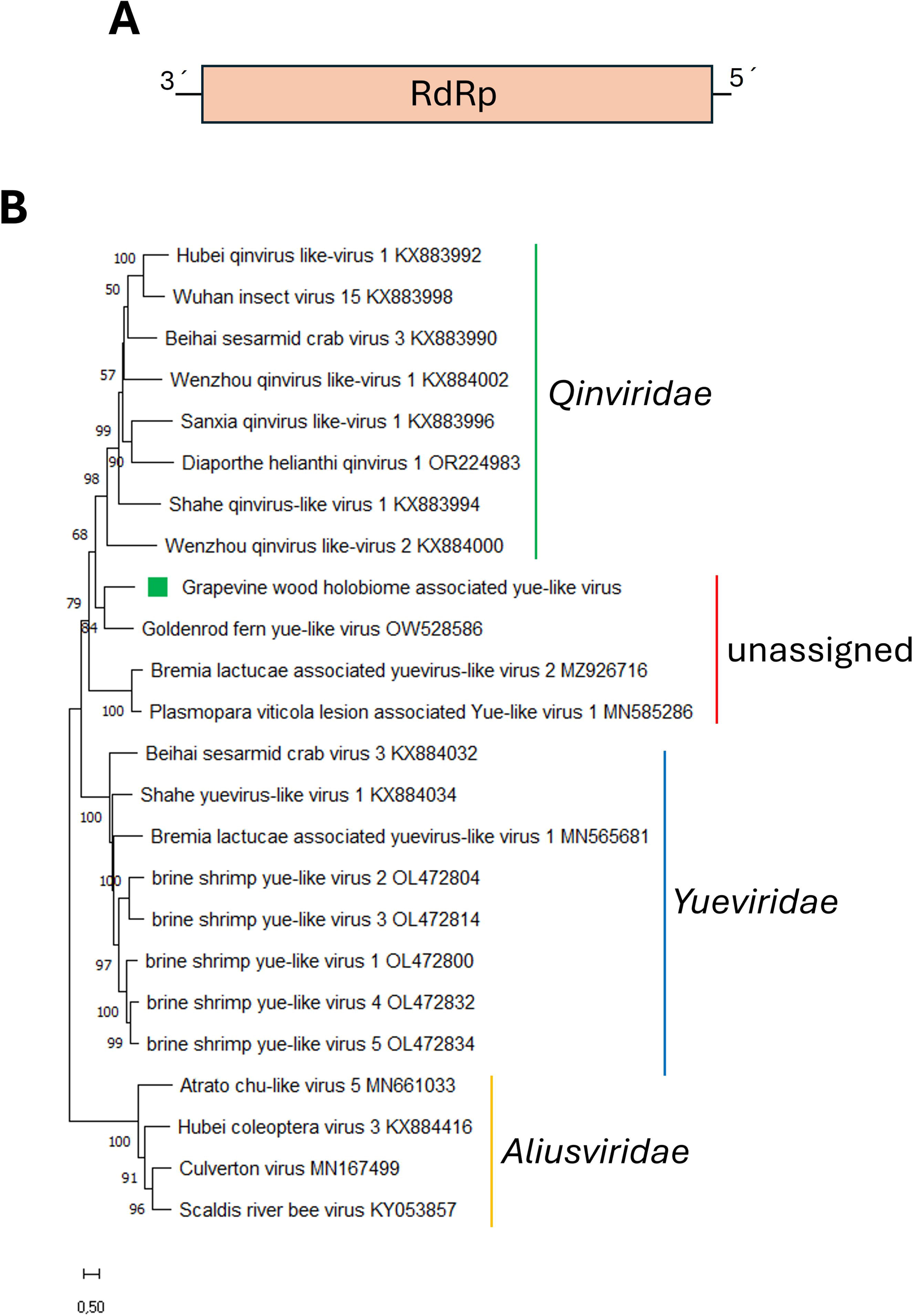
(**A**) Schematic representation of the genome organization of grapevine wood holobiome associated yue-like virus. (**B**) Maximum likelihood phylogenetic trees reconstructed using the RdRp protein sequence of grapevine wood holobiome associated yue-like virus and of representative yue-like and qin-like viruses. Bootstrap values above 50% are shown (1000 replicates). The grapevine wood holobiome associated yue-like virus is indicated with a green square. The scale bar shows the substitution per site.

A phylogenetic tree based on RdRp aa sequence alignment placed GWHaYLV within a well-supported clade with Goldenrod fern yue-like virus and other recently described yue-like viruses associated with fungal hosts (Fig. 14B). These viruses form a distinct clade separate from other previously classified viruses related to members of the *Qinviridae* family (Chiapello et al., 2020; Buivydaite et al., 2024; Forgia et al., 2024). All these viruses share a conserved IDD tripled in the Motif C of catalytic core of the RdRp (Buivydaite et al., 2024), and the same IDD motif was also identified in the RdRp of GWHaYKV1. Phylogenetic analysis and the presence of the IDD motif suggest that this group of viruses is linked to the *Qinviridae* family, which is a family of negative-sense RNA viruses with two-segmented genomes that have been associated with crustaceans, insects, gastropods, and nematodes (Wolf et al., 2023). Therefore, GWHaYLV represents a novel member of an unassigned virus family that belongs to the same order as *Qinviridae*. Finally, it is important to note that the GWHaYLV genome is likely incomplete, as no additional genome segments were identified, in contrast to what has been reported for Bremia lactucae associated Yue-like virus 2 and Plasmopara viticola associate Yuevirus1 (Chiapello et al., 2020; Forgia et al., 2024).

### 3.2 C. Double-stranded RNA viruses Botybirnaviruses

Two contigs showed similarity to sequences from members of the genus *Botybirnavirus*, whose members have a genome composed of two double-stranded RNA (dsRNA) segments encapsidated in isometric virions (Wu et al., 2012; Ye et al., 2023). This genus is the only one classified within the recently created family *Botybirnaviridae* (Simmonds et al., 2024). The putative botybirnavirus was named as grapevine wood holobiome associated botybirnavirus (GWHaBbV). The partial sequences of both RNA segments of this virus were obtained and deposited in GenBank under accession numbers PQ630621 and PQ630622 (Table 1). Each dsRNA contains a single ORF (Fig. 15A). The ORF of dsRNA1 encodes the CAP-POL fusion protein (Fig. 15A), with the RdRp domain identified in its C-terminal region, as reported in all botybirnaviruses to date (Ye et al., 2023). The N-terminal region likely contains a coat protein gene as previously described for botybirnaviruses (Ye et al., 2023). On the other hand, the ORF in dsRNA2 encodes a putative HP (Fig. 15A). This genomic organization is consistent with that reported for other botybirnaviruses (Cottet et al., 2019; Wu et al., 2012; Zhai et al., 2019; Ran et al., 2016; Xiang et al., 2017; Wang et al., 2018). BLASTp searches of the nr protein database using the CAP-POL protein and HP of GWHaBbV1 showed that the best hit were the RdRp and HP encoded by Leptosphaeria biglobosa botybirnavirus 2 (Table 1).

**Figure 15.**
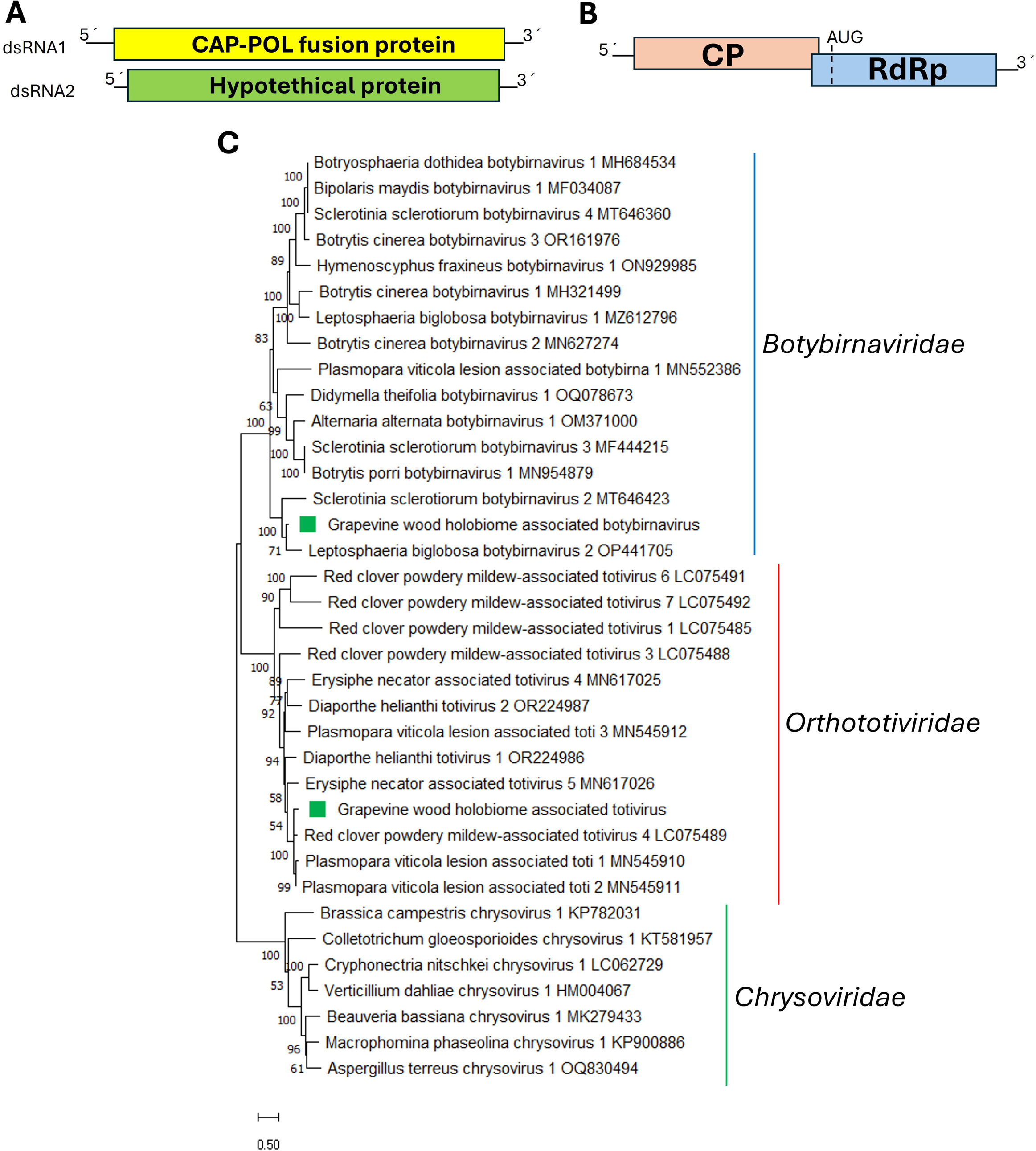
(**A**) Schematic representation of the genome organization of grapevine wood holobiome associated botybirnavirus. (**B**) Schematic representation of the genome organization of grapevine wood holobiome associated totivirus. (**C**) Maximum likelihood phylogenetic trees reconstructed using the RdRp protein sequence of grapevine wood holobiome associated botybirnavirus, grapevine wood holobiome associated totivirus, and of representative botybirnaviruses and totiviruses. Chrysovirids sequences were used as outgroup. Bootstrap values above 50% are shown (1000 replicates). The grapevine wood holobiome associated botybirnavirus and grapevine wood holobiome associated totivirus are indicated with a green square. The scale bar shows the substitution per site.

The phylogenetic tree based on RdRp aa sequence alignment placed GWHaBbV1 with other botyvirnaviruses, clustering with Leptosphaeria biglobosa botybirnavirus 2 and Sclerotinia sclerotiorum botybirnavirus 2 (Fig. 15C). Therefore, GWHaBbV represents a novel member of the genus *Botybirnavirus*.

Some botybirnaviruses have been reported to induce hypovirulence in their fungal hosts (Wu et al., 2012; Ran et al., 2016; Zhai et al., 2019; Liang et al., 2022). Therefore, further studies on the molecular characterization and the pathogenicity of GWHaBbV should be conducted to evaluate its potential as a biocontrol agent for fungal crop disease management.

#### Partitiviruses

Three contigs showed similarity to sequences from members of family *Partitiviridae,* whose members have a genome composed of two dsRNA segments (Vainio et al., 2018). The two identified segments correspond to two putative viruses, which were named grapevine wood holobiome associated partitivirus 1-2 (GWHaPV1-2). For GWHaPV1, the partial sequence of one segment was obtained and deposited in GenBank under accession number PV022023, while for GWHaPV2, two segments were assembled, one including the full-length coding region, while the other partially assembled, and deposited in GenBank under accession numbers PV0220234 and PV022025 (Table 1).

For GWHaPV1 the RNA segment encoding a putative RdRp was partially assembled (Fig. 16A). BLASTp searches of the nr protein database showed that the best hit was the RdRp protein encoded by Phomopsis vexans partitivirus 1 (Table 1). The full-length coding region of GWHaPV2 RNA1 is 1678 nt and encodes a putative RdRp (Fig. 16B) of 539 aa. BLASTp searches showed that the best hit was the RdRp protein encoded by Dothistroma septosporum partitivirus 1 (coverage, 99%; E-value, 0.0; identity, 75.46%). The partially assembled RNA2 encodes a putative CP (Fig. 16B), and the results of a BLASTp search showed that the best hit was the CP encoded by Plasmopara viticola lesion associated Partitivirus 3 (Table 1).

**Figure 16.**
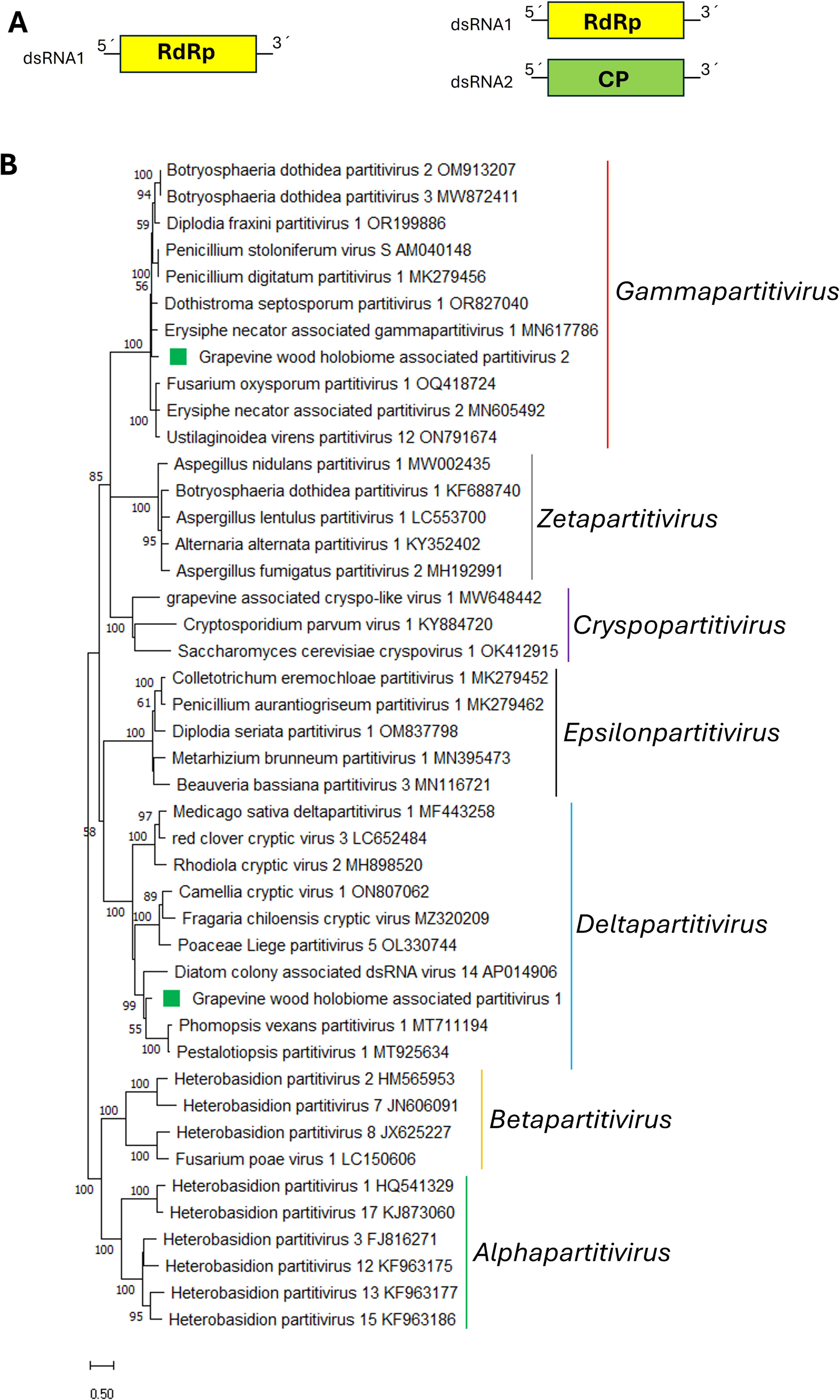
(**A**) Schematic representation of the genome organization of grapevine wood holobiome associated partitivirus 1. (**B**) Schematic representation of the genome organization of grapevine wood holobiome associated partitivirus 2. (**C**) Maximum likelihood phylogenetic trees reconstructed using the RdRp protein sequence of grapevine wood holobiome associated partitivirus 1-2 and of representative partitiviruses. Bootstrap values above 50% are shown (1000 replicates). The grapevine wood holobiome associated partitivirus 1-2 are indicated with a green square. The scale bar shows the substitution per site.

The phylogenetic tree based on RdRp aa sequence alignment placed GWHaPV1 with other deltapartitiviruses, clustering with Phomopsis vexans partitivirus 1 and Pestalotiopsis partititvirus 1 (Fig. 16C), while GWHaPV2 was placed in another cluster with other gammapartitiviruses (Fig. 16C).

#### Totiviruses

One contig showed similarity to sequences from members of the family *Orthototiviridae,* consisting of viruses with single linear dsRNA genomes that contain two, usually overlapping, ORFs encoding a putative CP and RdRp (Wickner et al., 2012). Only one genus, *Totivirus*, has been assigned within this family to accommodate fungal viruses (Simmonds et al., 2024). The threshold to demarcate species in this genus is an aa identity of 50% of their encoded proteins (Wickner et al., 2012). However, a great number of totiviruses have been identified in recent years (Xu et al., 2024; Chiapello et al., 2020); thus a 90% aa identity threshold may be more appropriate. The identified virus was named grapevine wood holobiome associated totivirus (GWHaTV). The coding-complete sequence of this virus was obtained and deposited in GenBank under accession number PQ630623 (Table 1).

The coding-complete sequence of GWHaTV is 4586 nt in length and contains two open reading frames (ORF1 and ORF2) (Fig. 15B). ORF1 encodes a CP of 683 aa, and a major capsid protein domain was identified in its sequence. The results of a BLASTp search of the nr protein database showed that the best hit was the CP encoded by Plasmopara viticola lesion associated totivirus 1 (coverage, 99%; E-value, 0.0; identity, 87.08%). ORF2 of GWHaTV encodes an RdRp of 806 aa, with the RdRp domain identified in its central region. The results of a BLASTp search showed that the best hit was the RdRp encoded by Plasmopara viticola lesion associated totivirus 1 (coverage, 100%; E-value, 0.0; identity, 79.53%). A putative slippery heptamer signal GGGUUUU, similar to that reported for Puccinia striiformis totiviruses (PsTVs) and red clover powdery mildew-associated totiviruses (Kondo et al., 2016; Zheng et al., 2017), was located 118 nt upstream of the ORF1 stop codon. This suggest that a -1 translational frameshifting strategy is involved in the expression of ORF2, as seen in other totiviruses, likely leading to the production of an ORF1–ORF2 (CP-RdRp) fusion protein (Khalifa and MacDiarmid, 2019) (Fig. 15B).

The phylogenetic tree based on RdRp aa sequence alignment placed GWHaATV with other totiviruses in a cluster with Plasmopara viticola lesion associated toti 1, Plasmopara viticola lesion associated toti 2 and red clover powdery mildew-associated totivirus 4 (Fig. 15C). Therefore, if the threshold of 90% is updated to demarcate species, GWHaTV is a novel member of the *Totiviridae* genus within the *Orthototiviridae* family.

### 3.3 Virus prevalence among samples

The prevalence of viruses among the analyzed grapevine samples exhibited substantial variability, with marked differences in detection levels and RNA read abundance across individual plants. A shifted detection and variable RNA levels among samples was observed (Table 2A-C), suggesting a complex dynamic in the persistence and expression of identified viruses. These differences likely reflect a combination of host-virus interactions, tissue tropism, and environmental influences affecting viral replication and accumulation.

#### Detection patterns of dsRNA and ss(-)RNA viruses

Analysis of dsRNA and ss(-)RNA viruses (Table 2A) revealed highly uneven distribution patterns among samples. For instance, GWHaBbV, GWHaBLV1, GWHaMOV1 and GWHaMNV1 were detected across all samples, albeit with significant variation in read counts, indicating their widespread presence in the grapevine holobiont. Notably, GWHaBLV1 exhibited particularly high read counts in symptomatic samples M16 (670/784) and M26 (393/506), suggesting a potential effect association with trunk disease. A striking case of selective presence was observed for instance with GWHaDV1, which was exclusively found in sample C25, where it reached exceptionally high read levels (1386/2144/2354). Similarly, GWHaMOV2 was abundant only in M11 (2485/2386), while being virtually undetectable in other samples (Table 2A). These findings highlight virus-host specificity, where certain viruses may be favored by particular physiological or pathological states of the plant. Moreover, despite low abundance in most samples, viruses such as GWHaTV and GWHaYLV displayed sporadic yet notable peaks, particularly in symptomatic plants M16 and M26, suggesting a potential effect of infection dynamics or secondary fungal associations.

#### Dominance of Ourmia-like, Mitoviruses, and Narna-like viruses

Ourmia-like, mitoviruses, and narna-like viruses were among the most prevalent virus groups detected (Table 2B). The exceptionally high read counts of GWHaOLV3, particularly in C22 (529,290) and M16 (77,599), indicate a dominant presence of this virus in these samples. Other notable cases include GWHaOLV2, which reached substantial levels in symptomatic M11 (2256) and M16 (1179), and GWHaOLV9, which showed consistently high levels across all samples. Mitoviruses were less uniformly distributed, with GWHaMV1 and GWHaMV5 detected in asymptomatic C22 and symptomatic M11, while GWHaMV3 was found exclusively in M26, reinforcing the idea of host-specific effect in its titters. Similarly, GWHaNLV1, a narna-like virus, exhibited high variability, ranging from undetectable levels in M11 to peak expression in M16 (3219). The high abundance of ourmia-like viruses in certain samples (e.g., GWHaOLV3 in C22) contrasts with their near absence in others, suggesting possible interactions with fungal hosts. This pattern aligns with previous reports that have identified these virus groups as the most abundant in fungal viromes associated with grapevine (Chiapello et al., 2020; Liu et al., 2023A).

#### Variable distribution of other ss(+)RNA viruses

The distribution of ss(+)RNA viruses (Table 2C) further supports the notion of dynamic virus-host interactions. GWHaAV1 and GWHaAV2 exhibited strong fluctuations in abundance, with GWHaAV1 reaching high levels in C25 (856) and M11 (514) but being absent in M26. Similarly, GWHaPLV2 showed an uneven pattern, being particularly abundant in C23 (3594) and C25 (2104), but with significantly reduced levels in M16 (77). Interestingly, GWHaDFV1, an ss(+)RNA virus, was almost exclusively found in C23 (9404), reinforcing the idea that certain viruses may be more compatible with specific plant physiological conditions or fungal endophytes. The absence of GWHaPLV1 in most samples except M26 (2260) suggests another case of localized viral dominance.

#### Implications for grapevine holobiont virome dynamics

The observed heterogeneity in virus prevalence suggests that the virome of grapevine wood is highly dynamic and influenced by multiple factors, including plant health status, microbial community interactions, and potential vector transmission. The significant presence of mitoviruses and narna-like viruses, which are typically associated with fungal hosts, further supports the hypothesis that many of these viruses persist within the grapevine mycobiome rather than in the plant itself. Moreover, the selective presence of certain viruses in symptomatic vs. asymptomatic plants raises important questions about their possible roles in disease progression or resistance. While some viruses, such as GWHaDV1, appear to be strongly associated with individual plants holobiomes, others, like GWHaOLV3, exhibit widespread prevalence, suggesting distinct ecological niches within the plant-fungi-virus interaction network. These findings highlight the complexity of viral prevalence patterns in the grapevine holobiont and underscore the need for further research to determine the functional implications of these viruses in plant health and disease. The high variability in RNA levels among samples (Table 2) suggests a non-uniform distribution of viral populations, likely shaped by host-virus compatibility, competition among fungal hosts, and environmental factors influencing viral replication.

## 4. Conclusions

The viral profile described here provides a first glimpse into the multifaceted South American grapevine wood holobiont virome. Through a comprehensive metatranscriptomic analysis of grapevine wood samples from both healthy plants and those exhibiting symptoms of grapevine trunk disease (GTD), we identified a striking diversity of viruses spanning multiple viral families and genomic architectures. Our findings reveal that mycoviruses associated with grapevine wood samples are abundant and taxonomically diverse, with representatives from established viral families such as *Botourmiaviridae*,

*Mitoviridae*, *Narnaviridae*, *Hypoviridae*, *Partitiviridae*, and *Discoviridae*, as well as from recently proposed families such as *Mycoaspiviridae* and *Splipalmiviridae*. Additionally, we report novel viruses with phylogenetic affinities to *Ambiguiviridae*, *Deltaflexiviridae*, *Bunyavirales*, *Picornavirales*, *Orthototiviridae*, and *Botybirnaviridae*, further expanding the known virosphere associated with grapevine fungal communities. Our findings hold important implications for understanding viral diversity, evolution, and ecological roles within the grapevine wood holobiont. The identification of highly divergent viruses suggests that novel genera—and in some cases, even new viral families—may need to be established to accommodate these newly discovered sequences. The classification of these viruses, particularly within the proposed *Mycoaspiviridae*, *Splipalmiviridae*, and novel clades of *Narnaviridae* and *Bunyavirales*, underscores the dynamic nature of virus evolution and the intricate host-virus interactions shaping fungal viromes in woody plant associated tissues. Beyond their taxonomic significance, some of the identified mycoviruses hold promise as potential biocontrol agents against GTD-associated fungal pathogens. Members of the families *Hypoviridae*, *Botourmiaviridae*, and *Botybirnaviridae* have been previously reported to induce hypovirulence in their fungal hosts, and our results suggest that similar mechanisms may be at play in certain newly identified viruses. Further functional characterization and host-range studies are required to determine whether these mycoviruses can be harnessed for sustainable disease management strategies in viticulture. Moreover, our study provides insights into the evolutionary trajectories of fungal-associated viruses and their potential interkingdom transmission events. The -presence of tombus-like, picorna-like, and bunya-like viruses—which are typically associated with plant and insect hosts—raises intriguing questions regarding viral movement across different biological compartments within the grapevine holobiont. This supports the growing body of evidence suggesting that viruses may transition between fungi, plants, and insect vectors more frequently than previously recognized. Taken together, these results significantly deepen our understanding of the diversity, taxonomy, and ecological roles of mycoviruses in the grapevine wood ecosystem. This work lays the foundation for future studies aimed at elucidating the functional impact of these viruses on fungal host dynamics, their potential interactions with other members of the grapevine microbiome, and their practical applications in disease management. The continued exploration of mycovirus diversity in perennial crops like grapevine may ultimately contribute to the development of innovative, virus-based biocontrol strategies to mitigate economically important plant diseases.

## Author contributions

**Humberto Debat**: data curation, formal analysis, writing-original draft, software, methodology, funding **Marcos Paolinelli**: biological data collection, data analysis, writing-review. **Georgina Escoriaza**: biological data collection, writing-review. **Sandra Garcia-Lampasona**: biological data collection, writing-review. **Sebastian Gomez-Talquenca**: biological data collection, writing-review, funding. **Nicolas Bejerman**: data curation, formal analysis, writing-review and editing, software, methodology, funding

## Acknowledgments

This work was partially supported by the Instituto Nacional de Tecnología Agropecuaria through the project INTA-PD-2023-I085.

## Data availability Statement

The raw data used to assembly the viruses described in the study is available in the ENA-EMBL repository [PRJEB31098 https://www.ebi.ac.uk/ena/data/view/PRJEB31098].

Viral sequences were deposited in the GenBank under accession numbers mentioned in Table 1 and the Manuscript.

## References

1. — Ayllón, M.A.; Turina, M.; Xie, J.; Nerva, L.; Marzano, S.-Y.L.; Donaire, L.; Jiang, D.; Consortium, I.R. ICTV Virus Taxonomy Profile: *Botourmiaviridae*. J. Gen. Virol. 2020, 101, 454–455.

2. — Ayllón, M.A.; Vainio, E.J. Chapter One—Mycoviruses as a Part of the Global Virome: Diversity, Evolutionary Links and Lifestyle. In Advances in Virus Research; Kielian, M., Roossinck, M.J., Eds.; Academic Press: Cambridge, MA, USA, 2023; Volume 115, pp. 1–86.

3. — Bass, D.; Stentiford, G.; Wang, H.-C.; Koskella, B.; Tyler, C. The Pathobiome in Animal and Plant Diseases. Trends Ecol. Evol. 2019, 34, 996–1008.

4. — Bankevich, A.; Nurk, S.; Antipov, D.; Gurevich, A.A.; Dvorkin, M.; Kulikov, A.S.; Lesin, V.M.; Nikolenko, S.I.; Pham, S.; Prjibelski, A.D.; et al. SPAdes: A new genome assembly algorithm and its applications to single-cell sequencing. J. Comput. Biol. 2012, 19, 455–477.

5. — Bettenfeld, P.; Cadena I Canals, J.; Jacquens, L.; Fernandez, O.; Fontaine, F.; van Schaik, E.; Courty, P.-E.; Trouvelot, S. The microbiota of the grapevine holobiont: A key component of plant health. J. Adv. Res. 2022, 40, 1–15.

6. — Bolger, A.M.; Lohse, M.; Usadel, B. Trimmomatic: A flexible trimmer for Illumina sequence data. Bioinformatics 2014, 30, 2114–2120.

7. — Botella, L.; Jung, M.H.; Rost, M.; Jung, T. Natural Populations from the Phytophthora Palustris Complex Show a High Diversity and Abundance of SsRNA and DsRNA Viruses. J. Fungi 2022, 8, 1118.

8. — Buchfink, B.; Xie, C.; Huson, D.H. Fast and sensitive protein alignment using DIAMOND. Nat. Methods 2015, 12, 59–60.

9. — Buivydaitė, Ž., Winding, A., Jørgensen, L. N., Zervas, A., Sapkota, R. New insights into RNA mycoviruses of fungal pathogens causing Fusarium head blight. Virus Res. 2024, 349, 199462.

10. — Charon, J.; Buchmann, J.P.; Sadiq, S.; Holmes, E.C. RdRp-scan: A bioinformatic resource to identify and annotate divergent RNA viruses in metagenomic sequence data. Virus Evol. 2022, 8, veac082.

11. — Chen, X.; He, H.; Yang, X.; Zeng, H.; Qiu, D.; Guo, L. The complete genome sequence of a novel *Fusarium graminearum* RNA virus in a new proposed family within the order Tymovirales. Arch. Virol. 2016, 161, 2899–2903.

12. — Chiapello, M.; Rodríguez-Romero, J.; Ayllón, M.A.; Turina, M. Analysis of the Virome Associated to Grapevine Downy Mildew Lesions Reveals New Mycovirus Lineages. Virus Evol. 2020, 6, veaa058.

13. — Chiba, Y.; Oiki, S.; Zhao, Y.; Nagano, Y.; Urayama, S.; Hagiwara, D. Splitting of RNA-Dependent RNA Polymerase Is Common in *Narnaviridae*: Identification of a Type II Divided RdRp from Deep-Sea Fungal Isolates. Virus Evol. 2021A, 7, veab095.

14. — Chiba, Y.; Oiki, S.; Yaguchi, T.; Urayama, S.; Hagiwara, D. Discovery of divided RdRp sequences and a hitherto unknown genomic complexity in fungal viruses. Virus Evol. 2021B, 7, veaa101.

15. — Chiba, S.; Velasco, L.; Ayllon, M.A.; Suzuki, N.; Lee-Marzano, S.Y.; Sun, L.; Sabanadzovic, S.; Turina, M. ICTV Virus Taxonomy Profile: *Hypoviridae* 2023. J. Gen. Virol. 2023, 104, 001848.

16. — Comont, G., Faure, C., Candresse, T., Laurens, M., Valière, S., Lluch, J., et al. Characterization of the RNA mycovirome associated with grapevine fungal pathogens: Analysis of mycovirus distribution and their genetic variability within a collection of Botryosphaeriaceae isolates. 2024, Viruses, 16, 392.

17. — Cottet, L., Potgieter, C. A., Castro, M. E., Castillo, A. Molecular characterization of a new botybirnavirus that infects Botrytis cinerea. Arch Virol, 2019, 164, 1479–1483.

18. — Dai, R., Yang, S., Pang, T., Tian, M., Wang, H., Zhang, D., et al. Identification of a negative-strand RNA virus with natural plant and fungal hosts. Proc Natl Acad Sci USA, 2024, 121(12), e2319582121.

19. — Daghino, S., Forgia, M., & Turina, M. Completion of the genome sequence of Oidiodendron maius splipalmivirus 1. Arch Virol, 2024, 169, 199.

20. — Dobin, A.; Gingeras, T.R. Mapping RNA-seq Reads with STAR. Curr. Protoc. Bioinform. 2015, *51*, 11.14.1–11.14.19.

21. — Donaire, L.; Rozas, J.; Ayllón, M.A. Molecular characterization of Botrytis ourmia-like virus, a mycovirus close to the plant pathogenic genus Ourmiavirus. Virology 2016, 489, 158–164.

22. — Fontaine, F.; Pinto, C.; Vallet, J.; Clément, C.; Gomes, A.C.; Spagnolo, A. The effects of grapevine trunk diseases (GTDs) on vine physiology. Eur. J. Plant Pathol. 2016, 144, 707–721.

23. — Forgia, M.; Daghino, S.; Chiapello, M.; Ciuffo, M.; Turina, M. New Clades of Viruses Infecting the Obligatory Biotroph Bremia Lactucae Representing Distinct Evolutionary Trajectory for Viruses Infecting Oomycetes. Virus Evol. 2024, 10, veae003.

24. — García-Pedrajas, M.D.; Cañizares, M.C.; Sarmiento-Villamil, J.L.; Jacquat, A.G.; Dambolena, J.S. Mycoviruses in Biological Control: From Basic Research to Field Implementation. Phytopathology 2019, 109, 1828–1839.

25. — Ghabrial, S.A.; Castón, J.R.; Jiang, D.; Nibert, M.L.; Suzuki, N. 50-plus Years of Fungal Viruses. Virology 2015, 479–480, 356–368.

26. — Gilbert, K.B.; Holcomb, E.E.; Allscheid, R.L.; Carrington, J.C. Hiding in plain sight: New virus genomes discovered via a systematic analysis of fungal public transcriptomes. PLoS ONE 2019, 14, e0219207.

27. — Grassi, F.; De Lorenzis, G. Back to the origins: Background and perspectives of grapevine domestication. Int. J. Mol. Sci. 2021, 22, 4518.

28. — Hamid, M.R.; Xie, J.; Wu, S.; Maria, S.K.; Zheng, D.; Assane Hamidou, A.; Wang, Q.; Cheng, J.; Fu, Y.; Jiang, D. A Novel Deltaflexivirus That Infects the Plant Fungal Pathogen, *Sclerotinia sclerotiorum*, Can Be Transmitted Among Host Vegetative Incompatible Strains. Viruses 2018, 10, 295.

29. — Hamim, I.; Urayama, S.i.; Netsu, O.; Tanaka, A.; Arie, T.; Moriyama, H.; Komatsu, K. Discovery, genomic sequence characterization and phylogenetic analysis of novel RNA viruses in the turfgrass pathogenic *Colletotrichum* spp. in Japan. Viruses 2022, 14, 2572.

30. — Hough, B.; Steenkamp, E.; Wingfield, B.; Read, D. Fungal viruses unveiled: A comprehensive review of mycoviruses. Viruses 2023, 15, 1202.

31. — Jacquat, A.G.; Theumer, M.G.; Dambolena, J.S. Putative Mitoviruses without In-Frame UGA(W) Codons: Evolutionary Implications. Viruses 2023, 15, 340.

32. — Jaillon, O.; Aury, J.M.; Noel, B.; Policriti, A.; Clepet, C.; Casagrande, A.; Choisne, N.; Aubourg, S.; Vitulo, N.; Vezzi, J.C.; et al. The grapevine genome sequence suggests ancestral hexaploidization in major angiosperm phyla. Nature 2007, 449, 463–467.

33. — Jia, J.; Fu, Y.; Jiang, D.; Mu, F.; Cheng, J.; Lin, Y.; Li, B.; Marzano, S.-Y.L.; Xie, J. Interannual Dynamics, Diversity and Evolution of the Virome in *Sclerotinia sclerotiorum* from a Single Crop Field. Virus Evol. 2021, 7, veab032.

34. — Jiāng, D.; Ayllón, M.A.; Marzano, S.-Y.L.; Kondō, H.; Turina, M. ICTV Virus Taxonomy Profile: *Mymonaviridae* 2022. J. Gen. Virol. 2022, 103, 001787.

35. — Jo, Y.; Choi, H.; Chu, H.; Cho, W.K. Unveiling Mycoviromes Using Fungal Transcriptomes. Int. J. Mol. Sci. 2022, 23, 10926.

36. — Katoh, K.; Standley, D.M. MAFFT multiple sequence alignment software version 7: Improvements in performance and usability. Mol. Biol. Evol. 2013, 30, 772–780.

37. — Kondo, H.; Hisano, S.; Chiba, S.; Maruyama, K.; Andika, I.B.; Toyoda, K.; Fujimori, F.; Suzuki, N. Sequence and phylogenetic analyses of novel totivirus-like double-stranded RNAs from field-collected powdery mildew fungi. Virus Res. 2016, 219, 353–364.

38. — Kondo, H.; Botella, L.; Suzuki, N. Mycovirus Diversity and Evolution Revealed/Inferred from Recent Studies. Annu. Rev. Phytopathol. 2022, 60, 307–336.

39. — Koonin, E.V. The Phylogeny of RNA-Dependent RNA Polymerases of Positive-Strand RNA Viruses. J. Gen. Virol. 1991, 72, 2197–2206.

40. — Kopylova, E.; Noé, L.; Touzet, H. SortMeRNA: Fast and accurate filtering of ribosomal RNAs in metatranscriptomic data. Bioinformatics 2012, 28, 3211–3217.

41. — Kotta-Loizou, I. Mycoviruses and Their Role in Fungal Pathogenesis. Curr. Opin. Microbiol. 2021, 63, 10–18.

42. — Kuhn, J., Adkins, S., Brown, K., De la Torre, J.C., Digiaro, M., et al. ICTV Virus Taxonomy Profile: Discoviridae 2023. J Gen Virol, 2023, 104, 001926.

43. — Kuhn, J.H.; Brown, K.; Adkins, S.; de la Torre, J.C.; Digiaro, M.; Ergunay, K.; Firth, A.E.; Hughes, H.R.; Junglen, S.; Lambert, A.J.; et al. Promotion of order Bunyavirales to class Bunyaviricetes to accommodate a rapidly increasing number of related polyploviricotine viruses. J. Virol. 2024, 98, e0106924.

44. — Le Gall, O.; Christian, P.; Fauquet, C.M.; King, A.M.; Knowles, N.J.; Nakashima, N.; Stanway, G.; Gorbalenya, A.E. Picornavirales, a proposed order of positive-sense single-stranded RNA viruses with a pseudo-T = 3 virion architecture. Arch. Virol. 2008, 153, 715–727.

45. — Leventhal, S.S.; Wilson, D.; Feldmann, H.; Hawman, D.W. A Look into Bunyavirales Genomes: Functions of Non-Structural (NS) Proteins. Viruses 2021, 13, 314.

46. — Li, K.; Zheng, D.; Cheng, J.; Chen, T.; Fu, Y.; Jiang, D.; Xie, J. Characterization of a novel *Sclerotinia sclerotiorum* RNA virus as the prototype of a new proposed family within the order Tymovirales. Virus Res. 2016, 219, 92–99.

47. — Li, H.; Bian, R.; Liu, Q.; Yang, L.; Pang, T.; Salaipeth, L.; Andika, I.B.; Kondo, H.; Sun, L. Identification of a novel hypovirulence-inducing hypovirus from *Alternaria alternata*. Front. Microbiol. 2019, 10, 1076.

48. — Li, Y.; Zhou, M.; Yang, Y.; Liu, Q.; Zhang, Z.; Han, C.; Wang, Y. Characterization of the mycovirome from the plant-pathogenic fungus Cercospora beticola. Viruses 2021, 13, 1915.

49. — Liang, Z.J.; Hua, H.H.; Wu, C.Y.; Zhou, T.; Wu, X.H. A botybirnavirus isolated from *Alternaria tenuissima* confers hypervirulence and decreased sensitivity of its host fungus to difenoconazole. Viruses 2022, 14, 2093.

50. — Liu, C., Jiang, X., Tan, Z., Wang, R., Shang, Q. Li, H. et al. An Outstandingly Rare Occurrence of Mycoviruses in Soil Strains of the Plant-Beneficial Fungi from the Genus Trichoderma and a Novel Polymycoviridae Isolate. 2023A, *Microbiol Spec*, *11*(3), e05228-22.

51. — Liu, H.Z.; Zhang, Y.F.; Liu, Y.Y.; Xiao, J.B.; Huang, Z.J.; Li, Y.F.; Li, H.P.; Li, P.F. Virome analysis of an ectomycorrhizal fungus *Suillus luteus* revealing potential evolutionary implications. Front. Cell. Infect. Microbiol. 2023B, 13, 1229859.

52. — Lozier, Z., Hill, L., Semmann, E., Miller, W. A. A proposed new Tombusviridae genus featuring extremely long 5’untranslated regions and a luteo/polerovirus-like gene block. Front Virol, 2024, 4, 1422934.

53. — Lu, X.; Dai, Z.; Xue, J.; Li, W.; Ni, P.; Xu, J.; Zhou, C.; Zhang, W. Discovery of Novel RNA Viruses through Analysis of Fungi-Associated next-Generation Sequencing Data. BMC Genom. 2024, 25, 517.

54. — Marais, A.; Faure, C.; Comont, G.; Candresse, T.; Stempien, E.; Corio-Costet, M.F. Characterization of the mycovirome of the phytopathogenic fungus, *Neofusicoccum parvum*. Viruses 2021, 13, 375.

55. — Marzano, S.-Y.L.; Domier, L.L. Novel mycoviruses discovered from metatranscriptomics survey of soybean phyllosphere phytobiomes. Virus Res. 2016, 213, 332–342.

56. — Mifsud, J.C.; Gallagher, R.V.; Holmes, E.C.; Geoghegan, J.L. Transcriptome Mining Expands Knowledge of RNA Viruses across the Plant Kingdom. J. Virol. 2022, 96, e00260–22.

57. — Morán, F., Olmos, A., Candresse, T., Ruiz-García, A. B. Complete Genome Characterization of Penicillimonavirus gammaplasmoparae, a Bipartite Member of the Family Mymonaviridae. Plants, 2023 12, 3300.

58. — Mu, F.; Li, B.; Cheng, S.; Jia, J.; Jiang, D.; Fu, Y.; Cheng, J.; Lin, Y.; Chen, T.; Xie, J. Nine viruses from eight lineages exhibiting new evolutionary modes that co-infect a hypovirulent phytopathogenic fungus. PLOS Pathog. 2021, 17, e1009823.

59. — Nerva, L.; Zanzotto, A.; Gardiman, M.; Gaiotti, F.; Chitarra, W. Soil microbiome analysis in an ESCA diseased vineyard. Soil Biol. Biochem. 2019A, 135, 60–70.

60. — Nerva, L.; Turina, M.; Zanzotto, A.; Gardiman, M.; Gaiotti, F.; Gambino, G.; Chitarra, W. Isolation, molecular characterization and virome analysis of culturable wood fungal endophytes in esca symptomatic and asymptomatic grapevine plants. Environ. Microbiol. 2019B, 21, 2886– 2904.

61. — Paolinelli, M.; Escoriaza, G.; Cesari, C.; Garcia-Lampasona, S.; Hernandez-Martinez, R. Characterization of Grapevine Wood Microbiome through a Metatranscriptomic Approach. Microb. Ecol. 2022, 83, 658–668.

62. — Ran, H.; Liu, L.; Li, B.; Cheng, J.; Fu, Y.; Jiang, D.; Xie, J. Co-infection of a hypovirulent isolate of *Sclerotinia sclerotiorum* with a new botybirnavirus and a strain of a mitovirus. Virol. J. 2016, 13, 92.

63. — Ruiz-Padilla, A.; Rodríguez-Romero, J.; Gómez-Cid, I.; Pacifico, D.; Ayllón, M.A. Novel Mycoviruses Discovered in the Mycovirome of a Necrotrophic Fungus. MBio 2021, 12, e03705–20.

64. — Sadiq, S.; Chen, Y.M.; Zhang, Y.Z.; Holmes, E.C. Resolving deep evolutionary relationships within the RNA virus phylum *Lenarviricota*. Virus Evol. 2022, 8, veac055.

65. — Sato, Y.; Shahi, S.; Telengech, P.; Hisano, S.; Cornejo, C.; Rigling, D.; Kondo, H.; Suzuki, N. A New Tetra-Segmented Splipalmivirus with Divided RdRP Domains from *Cryphonectria naterciae*, a Fungus Found on Chestnut and Cork Oak Trees in Europe. Virus Res. 2022, 307, 198606.

66. — Simmonds, P.; Adriaenssens, E.M.; Lefkowitz, E.J.; Oksanen, H.M.; Siddell, S.G.; Zerbini, F.M.; Alfenas-Zerbini, P.; Aylward, F.O.; Dempsey, D.M.; Dutilh, B.E.; et al. Changes to virus taxonomy and the ICTV Statutes ratified by the International Committee on Taxonomy of Viruses. Arch. Virol. 2024, 169, 236.

67. — Sutela, S.; Forgia, M.; Vainio, E.J.; Chiapello, M.; Daghino, S.; Vallino, M.; Martino, E.; Girlanda, M.; Perotto, S.; Turina, M. The Virome from a Collection of Endomycorrhizal Fungi Reveals New Viral Taxa with Unprecedented Genome Organization. Virus Evol. 2020, 6, veaa076.

68. — Tamura, K.; Stecher, G.; Kumar, S. MEGA11: Molecular Evolutionary Genetics Analysis Version 11. Mol. Biol. Evol. 2021, 38, 3022–3027.

69. — Theis, K.R.; Dheilly, N.M.; Klassen, J.L.; Brucker, R.M.; Baines, J.F.; Bosch, T.C.; Cryan, J.F.; Gilbert, S.F.; Goodnight, C.J.; Lloyd, E.A.; et al. Getting the Hologenome Concept Right: An Eco-Evolutionary Framework for Hosts and Their Microbiomes. mSystems 2016, 1, e00028–16.

70. — Urzo, M. L. R., Guinto, T. D., Eusebio-Cope, A., Budot, B. O., Yanoria, M. J. T., Jonson, G. B., et al. Metatranscriptomic sequencing of sheath blight-associated isolates of rhizoctonia solani revealed multi-infection by diverse groups of RNA viruses. 2024, Viruses, 16, 1152.

71. — Vainio, E.J.; Chiba, S.; Ghabrial, S.A.; Maiss, E.; Roossinck, M.; Sabanadzovic, S.; Suzuki, N.; Xie, J.; Nibert, M. ICTV virus taxonomy profile: Partitiviridae. J. Gen. Virol. 2018, 99, 17.

72. — Vayssier-Taussat, M.; Albina, E.; Citti, C.; Cosson, J.-F.; Jacques, M.-A.; Lebrun, M.; Le Loir, Y.; Ogliastro, M.; Petit, M.-A.; Roumagnac, P.; et al. Shifting the paradigm from pathogens to pathobiome: New concepts in the light of meta-omics. Front. Cell. Infect. Microbiol. 2014, 4, 29.

73. — Vazquez, A.L.; Alonso, J.M.M.; Parra, F. Mutation analysis of the GDD sequence motif of a calicivirus RNA-dependent RNA polymerase. J. Virol. 2000, 74, 3888–3891.

74. — Villan Larios, D.C.; Diaz Reyes, B.M.; Pirovani, C.P.; Loguercio, L.L.; Santos, V.C.; Góes-Neto, A.; Fonseca, P.L.C.; Aguiar, E.R.G.R. Exploring the mycovirus universe: Identification, diversity, and biotechnological applications. J. Fungi 2023, 9, 361.

75. — Wang, H.; Li, C.; Cai, L.; Fang, S.; Zheng, L.; Yan, F.; Zhang, S.; Liu, Y. The complete genomic sequence of a novel botybirnavirus isolated from a phytopathogenic *Bipolaris maydis*. Virus Genes 2018, 54, 733–736.

76. — Wang, Q.; Mu, F.; Xie, J.; Cheng, J.; Fu, Y.; Jiang, D. A single ssRNA segment encoding RdRp is sufficient for replication, infection, and transmission of ourmia-like virus in fungi. Front. Microbiol. 2020, 11, 379.

77. — Wang, Y.; Xu, Z.; Hai, D.; Huang, H.; Cheng, J.; Fu, Y.; Lin, Y.; Jiang, D.; Xie, J. Mycoviromic analysis unveils complex virus composition in a hypovirulent strain of *Sclerotinia sclerotiorum*. J. Fungi 2022, 8, 649.

78. — Wickner R. B.; Ghabrial S. A.; Nibert M. L.; Patterson J. L.; Wang C. C. 2012. “Family Totiviridae,” in Virus Taxonomy: Classification and Nomenclature of Viruses: Ninth Report of the International Committee on Taxonomy of Viruses, eds King A. M. Q., Adams M. J., Carstens E. B., Lefkowits E. J. (London: Elsevier Academic Press), 639–650.

79. — Wolf, Y.I.; Kazlauskas, D.; Iranzo, J.; Lucía-Sanz, A.; Kuhn, J.H.; Krupovic, M.; Dolja, V.V.; Koonin, E.V. Origins and Evolution of the Global RNA Virome. Mbio 2018, 9, e02329–18.

80. — Wolf, Y.I.; Koonin, E.V.; Krupovic, M.; Kuhn, J.H. ICTV Virus Taxonomy Profile: *Qinviridae* 2023. J. Gen. Virol. 2023, 104, 001905.

81. — Wu, M.; Jin, F.; Zhang, J.; Yang, L.; Jiang, D.; Li, G. Characterization of a Novel Bipartite Double-Stranded RNA Mycovirus Conferring Hypovirulence in the Phytopathogenic Fungus *Botrytis porri*. Virol. J 2012, 86, 6605–6619.

82. — Wu, C. F., Okada, R., Neri, U., Chang, Y. C., Ogawara, T., Kitaura, K., et al. Identification of a novel mycovirus belonging to the “flexivirus”-related family with icosahedral virion. 2024, Virus Evol, 10, veae093.

83. — Wu, C., Regedanz, E., Mathew, F., Kashyap, R., Mohan, K., Marzano, S-Y. L. Mycovirome of Diaporthe helianthi and D. gulyae, causal agents of Phomopsis stem canker of sunflower (Helianthus annuus L.). Virus Res. 2025, 351, 199521.

84. — Xiang, J.; Fu, M.; Hong, N.; Zhai, L.; Xiao, F.; Wang, G. Characterization of a novel botybirnavirus isolated from a phytopathogenic *Alternaria* fungus. Arch. Virol. 2017, 162, 3907– 3911.

85. — Xiao, J.B.; Wang, X.; Zheng, Z.R.; Wu, Y.G.; Wang, Z.; Li, H.P.; Li, P.F. Molecular characterization of a novel deltaflexivirus infecting the edible fungus *Pleurotus ostreatus*. Arch. Virol. 2023, 168, 162.

86. — Xie J., Jiang D. Understanding the diversity, evolution, ecology, and applications of mycoviruses. Annu. Rev. Microbiol. 2024, 78, 595–620.

87. — Xu, Z., Gao, Y. N., Teng, K., Ge, H., Zhang, X., Wu, M., et al. Identification and Genome Characterization of a Novel Virus within the Genus Totivirus from Chinese Bayberry (Myrica rubra). Viruses, 2024, 16, 283.

88. — Yao, Z.; Zou, C.; Peng, N.; Zhu, Y.; Bao, Y.; Zhou, Q.; Wu, Q.; Chen, B.; Zhang, M. Virome Identification and Characterization of Fusarium sacchari and F. andiyazi: Causative Agents of Pokkah Boeng Disease in Sugarcane. Front. Microbiol. 2020, 11, 240.

89. — Ye, L., Shi, X., He, Y., Chen, J., Xu, Q., Shafik, K., et al. A novel botybirnavirus with a unique satellite dsRNA causes latent infection in Didymella theifolia isolated from tea plants. Microbiol Spec, 2023, 11, e00033–23.

90. — Zell, R.; Groth, M.; Selinka, L.; Selinka, H.C. Picorna-like viruses of the Havel River, Germany. Front. Microbiol. 2022, 13, 865287.

91. — Zhai, L.; Yang, M.; Zhang, M.; Hong, N.; Wang, G. Characterization of a Botybirnavirus Conferring Hypovirulence in the Phytopathogenic Fungus *Botryosphaeria dothidea*. Viruses 2019, 11, 266.

92. — Zheng, L.; Lu, X.; Liang, X.; Jiang, S.; Zhao, J.; Zhan, G.; Liu, P.; Wu, J.; Kang, Z. Molecular characterization of novel Totivirus-like double-stranded RNAs from *Puccinia striiformis* f. sp.tritici, the causal agent of wheat stripe rust. Front. Microbiol. 2017, 8, 1960.

93. — Zhong, J.; Li, P.; Gao, B.D.; Zhong, S.Y.; Li, X.G.; Hu, Z.; Zhu, J.Z. Novel and diverse mycoviruses co-infecting a single strain of the phytopathogenic fungus *Alternaria dianthicola*. Front. Cell. Infect. Microbiol. 2022, 12, 980970.

