## Supplementary material for "Grapevine holobiome metatranscriptomics provides a glimpse into the wood mycovirome": Table 1

Table 1A. dsRNA and ss(-)RNA viruses identified

| Virus name | Virus acronym | Genome type | Segment number | Accesssion number | Contig lenght | Protein ID | Best match virus | Amino acid identity (%) |
| --- | --- | --- | --- | --- | --- | --- | --- | --- |
| grapevine wood holobiome associated botybirnavirus | GWHaBbV | dsRNA | 1  2 | PQ630621  PQ630622 | *  * | Cap-pol  HP | Leptosphaeria biglobosa botybirnavirus 2  Leptosphaeria biglobosa botybirnavirus 2 | *  * |
| grapevine wood holobiome associated totivirus | GWHaPV1 | dsRNA | 1 | PQ630623 | 4586 | CP  RdRp | Plasmopara viticola lesion associated toti 1 | 87.08  79.53 |
| grapevine wood holobiome associated partitivirus 1 | GWHaPV2 | dsRNA | 1 | PV022023 | * | RdRp | Phomopsis vexans partitivirus 1 | * |
| grapevine wood holobiome associated partitivirus 2 | GWHaTV | dsRNA | 1  2 | PV022024  PV022025 | 1678  * | RdRp  CP | Dothistroma septosporum partitivirus 1  Plasmopara viticola lesion associated Partitivirus 3 | 75.46  * |
| grapevine wood holobiome-associated bunya-like virus 1 | GWHaBLV1 | ss(-)RNA | 1  2 | PQ683342  PQ683343 | *  1016 | RdRp  NC | Sclerotinia sclerotiorum bunyavirus 6  Tulip streak virus | *  34.25 |
| grapevine wood holobiome-associated bunya-like virus 2 | GWHaBLV2 | ss(-)RNA | 1 | PQ683344 | * | RdRp | Plasmopara viticola lesion associated  mycobunyavirales-like virus 3 | * |
| grapevine wood holobiome-associated discovirus 1 | GWHaDV1 | ss(-)RNA | 1  2  3 | PQ508371  PQ508372  PQ508373 | 6502  1821  1195 | RdRp  NS  NC | Leptosphaeria biglobosa negative ssRNA virus 3  Leptosphaeria biglobosa negative ssRNA virus 3  Leptosphaeria biglobosa negative ssRNA virus 9 | 65.19  33.7  66.55 |
| grapevine wood holobiome-associated discovirus 2 | GWHaDV2 | ss(-)RNA | 1  2 | PQ508374  PQ508375 | 1563  1135 | NS  NC | Plasmopara viticola lesion associated  mycobunyavirales-like virus 4 | 30.68  79.65 |
| grapevine wood holobiome-associated mononegaambivirus1 | GWHaMNV1 | ss(-)RNA | 1  2 | PQ643870  PQ643871 | 6700  3651 | NC  RdRp  HP1  HP2  HP3  HP4 | Plasmopara viticola lesion associated mononegaambi virus 2  Plasmopara viticola lesion associated mononegaambi virus 2  Exserohilum turcicum mymonavirus 1  Plasmopara viticola lesion associated mononegaambi virus 2  Plasmopara viticola lesion associated mononegaambi virus 2  no hits | 38.67  69.65  40.23  66.24  59.39  - |
| grapevine wood holobiome-associated mononegaambivirus2 | GWHaMNV2 | ss(-)RNA | 1  2 | PQ643872  PQ643873 | 6777  3664 | NC  RdRp  HP1  HP2  HP3  HP4 | Plasmopara viticola lesion associated mononegaambi virus 2  Plasmopara viticola lesion associated mononegaambi virus 2  Exserohilum turcicum mymonavirus 1  Plasmopara viticola lesion associated mononegaambi virus 2  Plasmopara viticola lesion associated mononegaambi virus 2  no hits | 34.64  69.26  44.14  64.71  61.70  - |
| grapevine wood holobiome-associated mycoophiovirus 1 | GWHaMOV1 | ss(-)RNA | 1  2 | PQ638356  PQ638357 | 7696  2911 | RdRp  HP1  HP2  NC | Leptosphaeria biglobosa negative single-stranded RNA virus 7  no hits  Colletotrichum associated negative-sense  single-stranded RNA virus 2 | 83.38  -  47.55  69.69 |
| grapevine wood holobiome-associated mycoophiovirus 2 | GWHaMOV2 | ss(-)RNA | 1  2 | PQ638358  PQ638359 | 7651  2872 | RdRp  HP1  HP2  NC | Cladosporium cladosporioides negative-stranded RNA virus 1  no hits  Colletotrichum associated negative-sense  single-stranded RNA virus 2 | 69.19  -  47.79  69.81 |
| grapevine wood holobiome-associated Yue-like virus | GWHaYLV | ss(-)RNA | 1 | PQ720401 | * | RdRp | Goldenrod fern yue-like virus | * |

* partial sequence

Table 1B. Ourmia-like, mito and narna-like viruses identified

| Virus name | Virus acronym | Genome type | Segment number | Accesssion number | Contig lenght | Protein ID | Best match virus | Amino acid identity (%) |
| --- | --- | --- | --- | --- | --- | --- | --- | --- |
| grapevine wood holobiome-associated ourmia-like virus 1 | GWHaOLV1 | ss(+)RNA | 1 | PQ773447 | 2922 | RdRp | Plasmopara viticola lesion associated  ourmia-like virus 62 | 77.41 |
| grapevine wood holobiome-associated ourmia-like virus 2 | GWHaOLV2 | ss(+)RNA | 1 | PQ773448 | 2731 | RdRp | Lwood associated botourmia-like virus 25 | 62.76 |
| grapevine wood holobiome-associated ourmia-like virus 3 | GWHaOLV3 | ss(+)RNA | 1 | PQ773449 | 2726 | RdRp | Hangzhou botourmia-like virus 5 | 62.45 |
| grapevine wood holobiome-associated ourmia-like virus 4 | GWHaOLV4 | ss(+)RNA | 1 | PQ773450 | 2833 | RdRp | Guiyang botourmia-like virus 1 | 43.42 |
| grapevine wood holobiome-associated ourmia-like virus 5 | GWHaOLV5 | ss(+)RNA | 1 | PQ773451 | 2574 | RdRp | Hulumbuir Botou tick virus 8 | 78.72 |
| grapevine wood holobiome-associated ourmia-like virus 6 | GWHaOLV6 | ss(+)RNA | 1 | PQ773452 | 2442 | RdRp | Erysiphe necator associated ourmia-like virus 62 | 60.29 |
| grapevine wood holobiome-associated ourmia-like virus 7 | GWHaOLV7 | ss(+)RNA | 1 | PQ773453 | * | RdRp | Phomopsis vexans ourmia-like virus 1 | * |
| grapevine wood holobiome-associated ourmia-like virus 8 | GWHaOLV8 | ss(+)RNA | 1 | PQ773454 | * | RdRp | Yellow silver pine associated  botourmia-like virus 66 | * |
| grapevine wood holobiome-associated ourmia-like virus 9 | GWHaOLV9 | ss(+)RNA | 1 | PQ773455 | 2522 | RdRp | Sopagat virus | 62.1 |
| grapevine wood holobiome-associated ourmia-like virus 10 | GWHaOLV10 | ss(+)RNA | 1 | PQ773456 | * | RdRp | Plasmopara viticola lesion associated  ourmia-like virus 82 | * |
| grapevine wood holobiome-associated ourmia-like virus 11 | GWHaOLV11 | ss(+)RNA | 1 | PQ773457 | 2611 | RdRp | Hangzhou botourmia-like virus 3 | 77.83 |
| grapevine wood holobiome-associated ourmia-like virus 12 | GWHaOLV12 | ss(+)RNA | 1 | PQ773458 | 3011 | RdRp | Botourmiaviridae sp. | 72.54 |
| grapevine wood holobiome-associated ourmia-like virus 13 | GWHaOLV13 | ss(+)RNA | 1 | PQ773459 | * | RdRp | Sanya botourmia-like virus 4 | * |
| grapevine wood holobiome-associated ourmia-like virus 14 | GWHaOLV14 | ss(+)RNA | 1 | PQ773460 | 2698 | RdRp | Hangzhou botourmia-like virus 5 | 63.56 |
| grapevine wood holobiome-associated ourmia-like virus 15 | GWHaOLV15 | ss(+)RNA | 1 | PQ773461 | * | RdRp | Streptophyte associated botourmia-like virus 17 | * |
| grapevine wood holobiome-associated ourmia-like virus 16 | GWHaOLV16 | ss(+)RNA | 1 | PQ773462 | * | RdRp | Pestalotiopsis botourmiavirus 3 | * |
| grapevine wood holobiome-associated ourmia-like virus 17 | GWHaOLV17 | ss(+)RNA | 1 | PQ773463 | * | RdRp | Streptophyte associated botourmia-like virus 17 | * |
| grapevine wood holobiome-associated ourmia-like virus 18 | GWHaOLV18 | ss(+)RNA | 1 | PQ773464 | * | RdRp | Plasmopara viticola lesion associated  ourmia-like virus 62 | * |
| grapevine wood holobiome-associated ourmia-like virus 19 | GWHaOLV19 | ss(+)RNA | 1 | PQ773465 | * | RdRp | Sonacir virus | * |
| grapevine wood holobiome-associated ourmia-like virus 20 | GWHaOLV20 | ss(+)RNA | 1 | PQ773466 | 2351 | RdRp | Colletotrichum camelliae botourmiavirus 1 | 52.51 |
| grapevine wood holobiome-associated ourmia-like virus 21 | GWHaOLV21 | ss(+)RNA | 1 | PQ773467 | 2731 | RdRp | Botourmiaviridae sp. | 67.61 |
| grapevine wood holobiome-associated ourmia-like virus 22 | GWHaOLV22 | ss(+)RNA | 1 | PQ773468 | * | RdRp | Plasmopara viticola lesion associated  ourmia-like virus 62 | * |
| grapevine wood holobiome-associated ourmia-like virus 23 | GWHaOLV23 | ss(+)RNA | 1 | PQ773469 | 2488 | RdRp | Plasmopara viticola lesion associated  ourmia-like virus 22 | 72.95 |
| grapevine wood holobiome-associated ourmia-like virus 24 | GWHaOLV24 | ss(+)RNA | 1 | PQ773470 | 2579 | RdRp | Botrytis cinerea ourmia-like virus 14 | 74.69 |
| grapevine wood holobiome-associated mitovirus 1 | GWHaMV1 | ss(+)RNA | 1 | PQ729981 | * | RdRp | Rhizoctonia solani mitovirus 15 | * |
| grapevine wood holobiome-associated mitovirus 2 | GWHaMV2 | ss(+)RNA | 1 | PQ729982 | * | RdRp | Hainan mito-like virus 26 | * |
| grapevine wood holobiome-associated mitovirus 3 | GWHaMV3 | ss(+)RNA | 1 | PQ729983 | * | RdRp | Rhizoctonia cerealis duamitovirus | * |
| grapevine wood holobiome-associated mitovirus 4 | GWHaMV4 | ss(+)RNA | 1 | PQ729984 | * | RdRp | grapevine-associated mitovirus 2 | * |
| grapevine wood holobiome-associated mitovirus 5 | GWHaMV5 | ss(+)RNA | 1 | PQ729985 | * | RdRp | Sichuan mountain mitovirus 12 | * |
| grapevine wood holobiome-associated mitovirus 6 | GWHaMV6 | ss(+)RNA | 1 | PQ729986 | * | RdRp | Guangxi riberbank mitovirus 3 | * |
| grapevine wood holobiome-associated narna-like virus 1 | GWHaNLV1 | ss(+)RNA | 1 | PV021966 | 3178 | RdRp | Mbeech-associated narna-like virus 4 | 81.81 |
| grapevine wood holobiome-associated narna-like virus 2 | GWHaNLV2 | ss(+)RNA | 1 | PV021967 | 3155 | RdRp | Yellow silver pine associated narna-like virus 48 | 81.12 |
| grapevine wood holobiome-associated narna-like virus 3 | GWHaNLV3 | ss(+)RNA | 1 | PV021968 | 2488 | RdRp | Erysiphe necator associated narnavirus 16 | 75.07 |
| grapevine wood holobiome-associated narna-like virus 4 | GWHaNLV4 | ss(+)RNA | 1 | PV021969 | 3383 | RdRp | Qianjiang botourmia-like virus 41 | 43.78 |
| grapevine wood holobiome-associated narna-like virus 5 | GWHaNLV5 | ss(+)RNA | 1 | PV021970 | * | RdRp | Sopafab virus | * |
| grapevine wood holobiome-associated narna-like virus 6 | GWHaNLV6 | ss(+)RNA | 1  2  3  4 | PV021971  PV021972  PV021973  PV021974 | 3447  1921  1637  977 | RdRp  HP1  HP2  HP1  HP2  HP1 | Plasmopara viticola lesion associated narnavirus 32  Aspergillus creber narnavirus 1  Aspergillus creber narnavirus 1  Aspergillus creber narnavirus 1  no hits  no hits | 49.32  40.44  57.66  63.07  -  - |
| grapevine wood holobiome-associated narna-like virus 7 | GWHaNLV7 | ss(+)RNA | 1  2  3 | PV021975  PV021976  PV0211977 | 3411  1615  863 | RdRp  HP1  HP2  HP1 | Diaphorte gulyae narnavirus 2  no hits  Aspergillus creber narnavirus 1  Aspergillus creber narnavirus 1 | 82.7  -  32.91  28.07 |
| grapevine wood holobiome-associated narna-like virus 8 | GWHaNLV8 | ss(+)RNA | 1  2  3 | PV0211978  PV0211979  PV021980 | 3388  1608  779 | RdRp  HP1  HP2  HP1 | Ophiocordyceps sinensis narnavirus 1  Aspergillus creber narnavirus 1  Aspergillus creber narnavirus 1  Aspergillus creber narnavirus 1 | 48.23  36  29.54  29.41 |
| grapevine wood holobiome-associated narna-like virus 9 | GWHaNLV9 | ss(+)RNA | 1  2  3 | PV021981  PV021982  PV021983 | 3798  1598  827 | RdRp  HP1  HP2  HP1 | Erysiphe necator associated narnavirus 1  no hits  Aspergillus creber narnavirus 1  Aspergillus creber narnavirus 1 | 56.68  -  32.13  38.05 |
| grapevine wood holobiome-associated narna-like virus 10 | GWHaNLV10 | ss(+)RNA | 1  2  3 | PV021984  PV021985  PV021986 | 3430  1627  793 | RdRp  HP1  HP2  HP1 | Erysiphe necator associated narnavirus 1  no hits  Aspergillus creber narnavirus 1  Aspergillus creber narnavirus 1 | 55.69  -  34.25  38.57 |
| grapevine wood holobiome-associated narna-like virus 11 | GWHaNLV11 | ss(+)RNA | 1  2  3 | PV021987  PV021988  PV021989 | 3444  1666  791 | RdRp  HP1  HP2  HP1 | Erysiphe necator associated narnavirus 1  Aspergillus creber narnavirus 1  Aspergillus creber narnavirus 1  Aspergillus creber narnavirus 1 | 53.55  33.9  34.05  43.52 |
| grapevine wood holobiome-associated narna-like virus 12 | GWHaNLV12 | ss(+)RNA | 1  2  3 | PV021990  PV021991  PV021992 | 2074  2127  1276 | RdRp  RdRp  HP1  HP2 | Downy mildew lesion associated splipalmivirus 5  Leptosphaeria biglobosa narnavirus 4  Aspergillus flavus narnavirus 1  Aspergillus flavus narnavirus 1 | 85.43  67.29  66.2  32.8 |
| grapevine wood holobiome-associated narna-like virus 13 | GWHaNLV13 | ss(+)RNA | 1  2  3 | PV021993  PV021994  PV021995 | 2008  2141  1361 | RdRp  RdRp  HP1  HP2 | Leptosphaeria biglobosa narnavirus 5  Leptosphaeria biglobosa narnavirus 7  Downy mildew lesion associated splipalmivirus 44  Downy mildew lesion associated splipalmivirus 44 | 74.03  72.27  71.23  61.96 |
| grapevine wood holobiome-associated narna-like virus 14 | GWHaNLV14 | ss(+)RNA | 1  2  3 | PV021996  PV021997  PV021998 | 2034  2110  1350 | RdRp  RdRp  HP1  HP2 | Aspergillus fumigatus narnavirus 2  Aspergillus fumigatus narnavirus 2  Aspergillus fumigatus narnavirus 2  Downy mildew lesion associated splipalmivirus 6 | 72.7  65.44  54.25  41.94 |
| grapevine wood holobiome-associated narna-like virus 15 | GWHaNLV15 | ss(+)RNA | 1  2  3 | PV021999  PV021200  PV021201 | 2005  2148  1344 | RdRp  RdRp  HP1  HP2 | Aspergillus fumigatus narnavirus 2  Aspergillus fumigatus narnavirus 2  Aspergillus fumigatus narnavirus 2  Downy mildew lesion associated splipalmivirus 6 | 69.51  66.97  54.72  41.13 |
| grapevine wood holobiome-associated narna-like virus 16 | GWHaNLV16 | ss(+)RNA | 1  2  3 | PV021202  PV021203  PV021204 | 1997  2045  1342 | RdRp  RdRp  HP1  HP2 | Lwood associated narna-like virus  Downy mildew lesion associated splipalmivirus 6  Downy mildew lesion associated splipalmivirus 6 Downy mildew lesion associated splipalmivirus 6 | 77.19  69.95  59.73  42.42 |
| grapevine wood holobiome-associated narna-like virus 17 | GWHaNLV17 | ss(+)RNA | 1  2  3 | PV021205  PV021206  PV021207 | 2430  2205  950 | RdRp  RdRp  HP1 | Downy mildew lesion associated splipalmivirus 2  Dracophyllum associated narna-like virus 3  Downy mildew lesion associated splipalmivirus 1 | 48.72  47.90  34.83 |
| grapevine wood holobiome-associated narna-like virus 18 | GWHaNLV18 | ss(+)RNA | 1  2  3 | PV021208  PV021209  PV021210 | 2144  2228  1424 | RdRp  RdRp  HP1  HP2 | Yellow silver pine associated narna-like virus 29  Cryphonectria naterciae splipalmivirus 1  Cryphonectria naterciae splipalmivirus 1  Cryphonectria naterciae splipalmivirus 1 | 76.09  59.57  55.15  37.82 |
| grapevine wood holobiome-associated narna-like virus 19 | GWHaNLV19 | ss(+)RNA | 1  2  3 | PV021211  PV021212  PV021213 | 1937  2096  1370 | RdRp  RdRp  HP1  HP2 | Neofusicoccum parvum narnavirus 2  Downy mildew lesion associated splipalmivirus 44  Downy mildew lesion associated splipalmivirus 44  Downy mildew lesion associated splipalmivirus 44 | 80.79  85.52  67.11  56.55 |
| grapevine wood holobiome-associated narna-like virus 20 | GWHaNLV20 | ss(+)RNA | 1  2  3 | PV021214  PV021215  PV021216 | 2372  2225  1118 | RdRp  RdRp  HP1 | Cladosporium tenuissimum narnavirus 1  Leptosphaeria biglobosa narnavirus 2  Sclerotinia sclerotiorum narnavirus 2 | 89.39  62.79  57.28 |
| grapevine wood holobiome-associated narna-like virus 21 | GWHaNLV21 | ss(+)RNA | 1  2  3 | PV021217  PV021218  PV021219 | 2330  2264  1145 | RdRp  RdRp  HP1 | Exserohilum turcicum narnavirus 1  Exserohilum turcicum narnavirus 1  Exserohilum turcicum narnavirus 1 | 86.23  80.93  58.01 |
| grapevine wood holobiome-associated narna-like virus 22 | GWHaNLV22 | ss(+)RNA | 1  2  3 | PV021220  PV021221  PV021222 | 2314  2303  1152 | RdRp  RdRp  HP1 | Exserohilum turcicum narnavirus 1  Exserohilum turcicum narnavirus 1  Exserohilum turcicum narnavirus 1 | 82.65  81.21  70.20 |

* partial sequence

Table 1C. other ss(+)RNA viruses identified

| Virus name | Virus acronym | Genome type | Segment number | Accesssion number | Contig lenght | Protein ID | Best match virus | Amino acid identity (%) |
| --- | --- | --- | --- | --- | --- | --- | --- | --- |
| grapevine wood holobiome associated ambiguivirus 1 | GWHaAV1 | ss(+)RNA | 1 | PQ474301 | 3214 | HP  RdRp | rice tombus-like virus 3  Alternaria dianthicola umbra-like virus 1 | 44.27  55.82 |
| grapevine wood holobiome associated ambiguivirus 2 | GWHaAV2 | ss(+)RNA | 1 | PQ474302 | 3201 | HP  RdRp | Erysiphe necator associated ambiguivirus 2  Alternaria dianthicola umbra-like virus 1 | 40.61  54.97 |
| grapevine wood holobiome associated ambiguivirus 3 | GWHaAV3 | ss(+)RNA | 1 | PQ590144 | 4096 | HP  RdRp | soybean leaf-associated ssRNA virus 1  grapevine-associated tombus-like virus 4 | 42.08  54.77 |
| grapevine wood holobiome-associated deltaflexivirus 1 | GWHaDV1 | ss(+)RNA | 1 | PQ567243 | 8592 | RdRp  HP1  HP2  HP3  HP4 | Fusarium deltaflexivirus 2  Guangxi alphaflexi-like virus  no hits  no hits  Sesame deltaflexivirus 1 | 58.51  31.25  -  -  27.44 |
| grapevine wood holobiome-associated deltaflexivirus 2 | GWHaDV2 | ss(+)RNA | 1 | PQ567244 | * | RdRp  HP1  HP2  HP3 | Erysiphe necator associated deltaflexivirus 3  no hits  Sclerotinia sclerotiorum deltaflexivirus 1  Sclerotinia sclerotiorum deltaflexivirus 1 | *  -  46.24  45.07 |
| grapevine wood holobiome-associated deltaflexivirus 3 | GWHaDV3 | ss(+)RNA | 1 | PQ567245 | * | RdRp | Pestalotiopsis deltaflexivirus 1 | * |
| grapevine wood holobiome-associated hypovirus | GWHaHV | ss(+)RNA | 1 | PQ673554 | 13684 | RdRp | Beticola hypovirus 1 | 56.21 |
| grapevine wood holobiome-associated picorna-like virus 1 | GWHaPLV1 | ss(+)RNA | 1 | PQ762193 | 9896 | RdRp | Picornavirales sp. | 43.07 |
| grapevine wood holobiome-associated picorna-like virus 2 | GWHaPLV2 | ss(+)RNA | 1 | PQ762194 | 9344 | CP  RdRp | Riboviria sp.  Riboviria sp. | 64.34  44.79 |
| grapevine wood holobiome-associated picorna-like virus 3 | GWHaPLV3 | ss(+)RNA | 1 | PQ762195 | * | RdRp | Riboviria sp. | * |
| grapevine wood holobiome-associated picorna-like virus 4 | GWHaPLV4 | ss(+)RNA | 1 | PQ762196 | 9158 | CP  RdRp | Riboviria sp.  Riboviria sp. | 62.84  54.93 |
| grapevine wood holobiome-associated picorna-like virus 5 | GWHaPLV5 | ss(+)RNA | 1 | PQ762197 | * | CP  RdRp | Dicistroviridae sp.  Dicistroviridae sp. | 51.34  * |
| grapevine wood holobiome-associated tombus-like virus | GWHaTLV | ss(+)RNA | 1 | PV036960 | * | HP  RdRp  HP  CP  HP | Chrocosarb virus  Erysiphe necator associated tombus-like virus 2  Nanning Tombu tick virus 1  Zizania latifolia tombusvirus  Zizania latifolia tombusvirus | *  86.65  33.5  90.14  75.82 |

* partial sequence
