## Supplementary material for "Grapevine holobiome metatranscriptomics provides a glimpse into the wood mycovirome": Table 2

Table 2A. dsRNA and ss(-) RNA virus presence among samples assessed by read mapping of individual libraries to consensus genomes/genome segments. Numbers indicate the absolute value of virus reads

|  |  | Library | | | | | |
| --- | --- | --- | --- | --- | --- | --- | --- |
|  |  | C22 | C23 | C25 | M11 | M16 | M26 |
| Virus | GWHaBbV | 15/16 | 129/112 | 27/51 | 126/108 | 48/55 | 65/46 |
|  | GWHaPV1 | 0 | 0 | 96 | 0 | 0 | 0 |
|  | GWHaPV2 | 75/25 | 2/4 | 214/103 | 0/2 | 0/5 | 2/7 |
|  | GWHaTV | 4 | 22 | 322 | 10 | 104 | 28 |
|  | GWHaBLV1 | 302/397 | 114/208 | 150/267 | 81/196 | 670/784 | 393/506 |
|  | GWHaBLV2 | 0 | 4 | 201 | 11 | 23 | 2 |
|  | GWHaDV1 | 0 | 0 | 1386/2144/2354 | 0 | 0 | 4/0/0 |
|  | GWHaDV2 | 96/10 | 10/38 | 6/36 | 16/20 | 4/22 | 54/52 |
|  | GWHaMNV1 | 242/115 | 3167/2968 | 24/15 | 36/21 | 3309/3174 | 632/435 |
|  | GWHaMNV2 | 0 | 2/0 | 12/3 | 758/576 | 2660/2489 | 0 |
|  | GWHaMOV1 | 232/114 | 160/96 | 2457/2348 | 833/695 | 40/24 | 210/103 |
|  | GWHaMOV2 | 0 | 20/12 | 0 | 2485/2386 | 6/3 | 0 |
|  | GWHaYLV | 2 | 4 | 0 | 0 | 108 | 36 |

*Virus name is shown in Table 1

Table 2B. Ourmia-like, mito and narna-like virus presence among samples assessed by read mapping of individual libraries to consensus genomes/genome segments. Numbers indicate the absolute value of virus reads

|  |  | Library | | | | | |
| --- | --- | --- | --- | --- | --- | --- | --- |
|  |  | C22 | C23 | C25 | M11 | M16 | M26 |
| Virus | GWHaOLV1 | 20 | 0 | 0 | 0 | 2699 | 20 |
|  | GWHaOLV2 | 76 | 192 | 124 | 2256 | 1179 | 404 |
|  | GWHaOLV3 | 529290 | 46464 | 12087 | 19054 | 77599 | 1315 |
|  | GWHaOLV4 | 0 | 0 | 3237 | 0 | 0 | 2 |
|  | GWHaOLV5 | 29 | 2126 | 0 | 0 | 2043 | 2 |
|  | GWHaOLV6 | 0 | 0 | 0 | 0 | 405 | 0 |
|  | GWHaOLV7 | 0 | 0 | 0 | 0 | 546 | 0 |
|  | GWHaOLV8 | 114 | 4 | 0 | 8 | 22 | 2 |
|  | GWHaOLV9 | 2190 | 288 | 622 | 1442 | 403 | 2302 |
|  | GWHaOLV10 | 18 | 90 | 129 | 62 | 97 | 70 |
|  | GWHaOLV11 | 1 | 0 | 14 | 0 | 186 | 4295 |
|  | GWHaOLV12 | 0 | 10 | 0 | 1036 | 2 | 1872 |
|  | GWHaOLV13 | 0 | 8 | 2 | 2405 | 0 | 315 |
|  | GWHaOLV14 | 408 | 3 | 0 | 5 | 0 | 0 |
|  | GWHaOLV15 | 139 | 227 | 1252 | 4 | 1 | 155 |
|  | GWHaOLV16 | 4 | 2 | 0 | 282 | 10 | 5 |
|  | GWHaOLV17 | 9 | 494 | 47 | 0 | 5 | 12 |
|  | GWHaOLV18 | 0 | 371 | 2 | 4 | 4 | 22 |
|  | GWHaOLV19 | 8 | 42 | 0 | 0 | 11 | 71 |
|  | GWHaOLV20 | 0 | 50 | 434 | 0 | 28 | 2 |
|  | GWHaOLV21 | 0 | 2 | 46 | 1782 | 156 | 64 |
|  | GWHaOLV22 | 0 | 4 | 1 | 188 | 0 | 36 |
|  | GWHaOLV23 | 0 | 2 | 0 | 636 | 0 | 0 |
|  | GWHaOLV24 | 0 | 2 | 0 | 893 | 6 | 8 |
|  | GWHaMV1 | 86 | 2 | 0 | 0 | 0 | 26 |
|  | GWHaMV2 | 0 | 0 | 36 | 0 | 0 | 82 |
|  | GWHaMV3 | 0 | 0 | 0 | 0 | 0 | 151 |
|  | GWHaMV4 | 205 | 0 | 0 | 4 | 0 | 10 |
|  | GWHaMV5 | 272 | 2 | 6 | 100 | 0 | 0 |
|  | GWHaMV6 | 18 | 0 | 0 | 72 | 0 | 0 |
|  | GWHaNLV1 | 311 | 2 | 1453 | 3 | 3219 | 6 |
|  | GWHaNLV2 | 745 | 72 | 44 | 116 | 66 | 240 |
|  | GWHaNLV3 | 105 | 143 | 457 | 14 | 10 | 328 |
|  | GWHaNLV4 | 0 | 0 | 0 | 0 | 0 | 754 |
|  | GWHaNLV5 | 32 | 23 | 0 | 150 | 0 | 0 |
|  | GWHaNLV6 | 0 | 0 | 0 | 0 | 1045/854/627/432 | 0 |
|  | GWHaNLV7 | 6/2/2 | 114/49/18 | 160/64/27 | 30/19/8 | 3856/2563/1866 | 21/11/5 |
|  | GWHaNLV8 | 40/23/9 | 0 | 4768/3218/2147 | 2/0/0 | 266/128/59 | 4245/2986/2043 |
|  | GWHaNLV9 | 258/169/87 | 191/118/56 | 110/68/23 | 2394/1684/1165 | 200/124/67 | 36/19/8 |
|  | GWHaNLV10 | 21/10/4 | 1/0/0 | 0 | 4/0/0 | 0 | 329/215/146 |
|  | GWHaNLV11 | 0 | 0 | 0 | 489/296/214 | 0 | 2/0/0 |
|  | GWHaNLV12 | 38/16/11 | 92/56/29 | 0 | 8/4/2 | 539/415/314 | 0 |
|  | GWHaNLV13 | 0 | 22/14/9 | 20/12/6 | 231/134/78 | 311/186/109 | 2/0/0 |
|  | GWHaNLV14 | 0 | 1/0/0 | 106/68/39 | 3/0/0 | 0 | 686/487/354 |
|  | GWHaNLV15 | 0 | 1/0/0 | 0 | 0 | 0 | 185/119/62 |
|  | GWHaNLV16 | 53/31/12 | 0 | 764/629/537 | 0 | 0 | 0 |
|  | GWHaNLV17 | 0 | 101/68/39 | 0 | 145/84/53 | 609/515/427 | 0 |
|  | GWHaNLV18 | 43/21/12 | 845/711/634 | 0 | 0 | 0 | 169/96/67 |
|  | GWHaNLV19 | 0 | 967/851/725 | 125/58/39 | 0 | 0 | 0 |
|  | GWHaNLV20 | 0 | 0 | 0 | 87/49/31 | 795/627/548 | 0 |
|  | GWHaNLV21 | 65/34/19 | 0 | 98/44/23 | 469/384/296 | 215/147/83 | 0 |
|  | GWHaNLV22 | 0 | 732/586/496 | 0 | 0 | 0 | 136/67/45 |

*Virus name is shown in Table 1

Table 2C. Other ss(+)RNA virus presence among samples assessed by read mapping of individual libraries to consensus genomes/genome segments. Numbers indicate the absolute value of virus reads

|  |  | Library | | | | | |
| --- | --- | --- | --- | --- | --- | --- | --- |
|  |  | C22 | C23 | C25 | M11 | M16 | M26 |
| Virus | GWHaAV1 | 148 | 8 | 856 | 514 | 146 | 0 |
|  | GWHaAV2 | 157 | 0 | 0 | 1425 | 14 | 0 |
|  | GWHaAV3 | 2 | 0 | 0 | 0 | 846 | 0 |
|  | GWHaDFV1 | 0 | 9404 | 96 | 0 | 14 | 0 |
|  | GWHaDFV2 | 0 | 0 | 0 | 522 | 0 | 0 |
|  | GWHaDFV3 | 0 | 0 | 0 | 0 | 0 | 102 |
|  | GWHaHV | 723 | 0 | 0 | 0 | 0 | 0 |
|  | GWHaPLV1 | 0 | 0 | 0 | 0 | 0 | 2260 |
|  | GWHaPLV2 | 731 | 3594 | 2104 | 1887 | 77 | 1447 |
|  | GWHaPLV3 | 0 | 0 | 0 | 2 | 0 | 394 |
|  | GWHaPLV4 | 1614 | 1384 | 399 | 2846 | 18 | 457 |
|  | GWHaPLV5 | 10 | 62 | 588 | 6 | 83 | 2 |
|  | GWHaTLV | 2 | 2 | 727 | 146 | 0 | 190 |

*Virus name is shown in Table 1

.
